# Structural basis of RNA-guided transcription by a dCas12f-σ^E^-RNAP complex

**DOI:** 10.1101/2025.06.10.658880

**Authors:** Renjian Xiao, Florian T. Hoffmann, Dan Xie, Tanner Wiegand, Adriana I. Palmieri, Samuel H. Sternberg, Leifu Chang

## Abstract

RNA-guided proteins have emerged as critical transcriptional regulators in both natural and engineered biological systems by modulating RNA polymerase (RNAP) and its associated factors^1–3^. In bacteria, diverse clades of repurposed TnpB and CRISPR-associated proteins repress gene expression by blocking transcription initiation or elongation, enabling non-canonical modes of regulatory control and adaptive immunity^1,4,5^. Intriguingly, a distinct class of nuclease-dead Cas12f homologs (dCas12f) instead activates gene expression through its association with unique extracytoplasmic function sigma factors (σ^E^)^6^, though the molecular basis has remained elusive. Here we reveal a novel mode of RNA-guided transcription initiation by determining cryo-electron microscopy structures of the dCas12f-σ^E^ system from *Flagellimonas taeanensis*. We captured multiple conformational and compositional states, including the DNA-bound dCas12f-σ^E^-RNAP holoenzyme complex, revealing how RNA-guided DNA binding leads to σ^E^-RNAP recruitment and nascent mRNA synthesis at a precisely defined distance downstream of the R-loop. Rather than following the classical paradigm of σ^E^-dependent promoter recognition, these studies show that recognition of the −35 element is largely supplanted by CRISPR-Cas targeting, while the melted −10 element is stabilized through unusual stacking interactions rather than insertion into the typical recognition pocket. Collectively, this work provides high-resolution insights into an unexpected mechanism of RNA-guided transcription, expanding our understanding of bacterial gene regulation and opening new avenues for programmable transcriptional control.

## INTRODUCTION

Bacterial transcription is catalyzed by a conserved RNA polymerase (RNAP) core enzyme comprising four distinct subunits: α, β, β’, and ω. RNAP exhibits only non-specific interactions with DNA^7^, whereas sequence-specific binding of promoter motifs during transcription initiation relies critically on a dedicated sigma (σ) factor that comprises a key part of the σ-RNAP holoenzyme complex^8^. Bacteria typically encode one RNAP core enzyme but multiple σ factors, each of which recognizes a distinct promoter sequence, thus allowing adaptation to diverse cellular conditions^9^. The primary Group 1 σ factor in *E. coli*, σ^70^, is responsible for transcribing essential housekeeping genes and contains four conserved domains (σ1–σ4), with σ4 binding the –35 element and σ2 recognizing the melted –10 element in standard *E. coli* σ^70^ promoters^8–10^. Group 2 and 3 σ factors play more specialized regulatory roles, including controlling genes involved in stationary phase, flagellar biosynthesis, and heat shock, and share many of the same conserved regions as σ^70^. Group 4 σ factors, also known as extracytoplasmic function (ECF) σ^E^, are structurally more compact and retain only the σ_2_ and σ_4_ domains connected by a σ finger (also known as σ_2/4_ linker)^11,12^ . Intriguingly, whereas *E. coli* encodes only a single homolog, σ^E^ factors are the largest and most diverse group in other bacterial taxa^13^, where they canonically respond to diverse environmental cues via interactions with their cognate anti-sigma factors that localize to the cellular membrane^13,14^.

Recent genomic analyses uncovered a subset of ECF σ^E^ factors encoded adjacent to atypical Cas12f-like homologs that lack the expected association with CRISPR arrays^15^. These unusual σ^E^ proteins feature a C-terminal extension resembling a helix-turn-helix (HTH) domain predicted to interact with Cas12f, and the presence of conserved Cas12f mutations in the catalytic RuvC domain, as well as the absence of C-terminal regions critical for DNA cleavage, further suggesting a novel function beyond adaptive immunity^15^. In particular, the strong co-occurrence between σ^E^ and nuclease-dead Cas12f (dCas12f) implied the possibility that guide RNA (gRNA) molecules might be involved in directing a new mode of transcriptional activation through direct interactions with RNAP. Alongside recently described systems that employ repurposed Cas9, Cas12, or TnpB-like homologs for transcriptional repression^1,4,5,16,17^, a natural mechanism of RNA-guided transcription initiation would expand the known roles of programmable gene regulation while also complementing existing technologies for CRISPR interference (CRISPRi) and CRISPR activation (CRISPRa) applications^2,18^.

Here, we present the biochemical and structural basis of RNA-guided transcription initiation by the dCas12f-σ^E^, whose cellular activities are detailed in a companion article^6^. Focusing on homologs from *Flagellimonas taeanensis* that regulate SusC/SusD-family outer membrane proteins^6^ (**Fig. 1a,b**), we resolved multiple cryo-EM structures including dCas12f-gRNA, DNA-bound dCas12f-gRNA, and the DNA-bound dCas12f-σ^E^-RNAP holoenzyme complex. Our structures elegantly reveal how recognition of a simple 5’-G target-adjacent motif (TAM) leads to R-loop formation, **σ**^E^-RNAP recruitment, and a tightly defined transcription start site (TSS) ∼46-bp downstream, without obligate recognition of typical promoter elements. These findings provide new insights into diverse σ factor function and inform future strategies for synthetic, promoter-independent gene activation tools.

**Figure 1.**
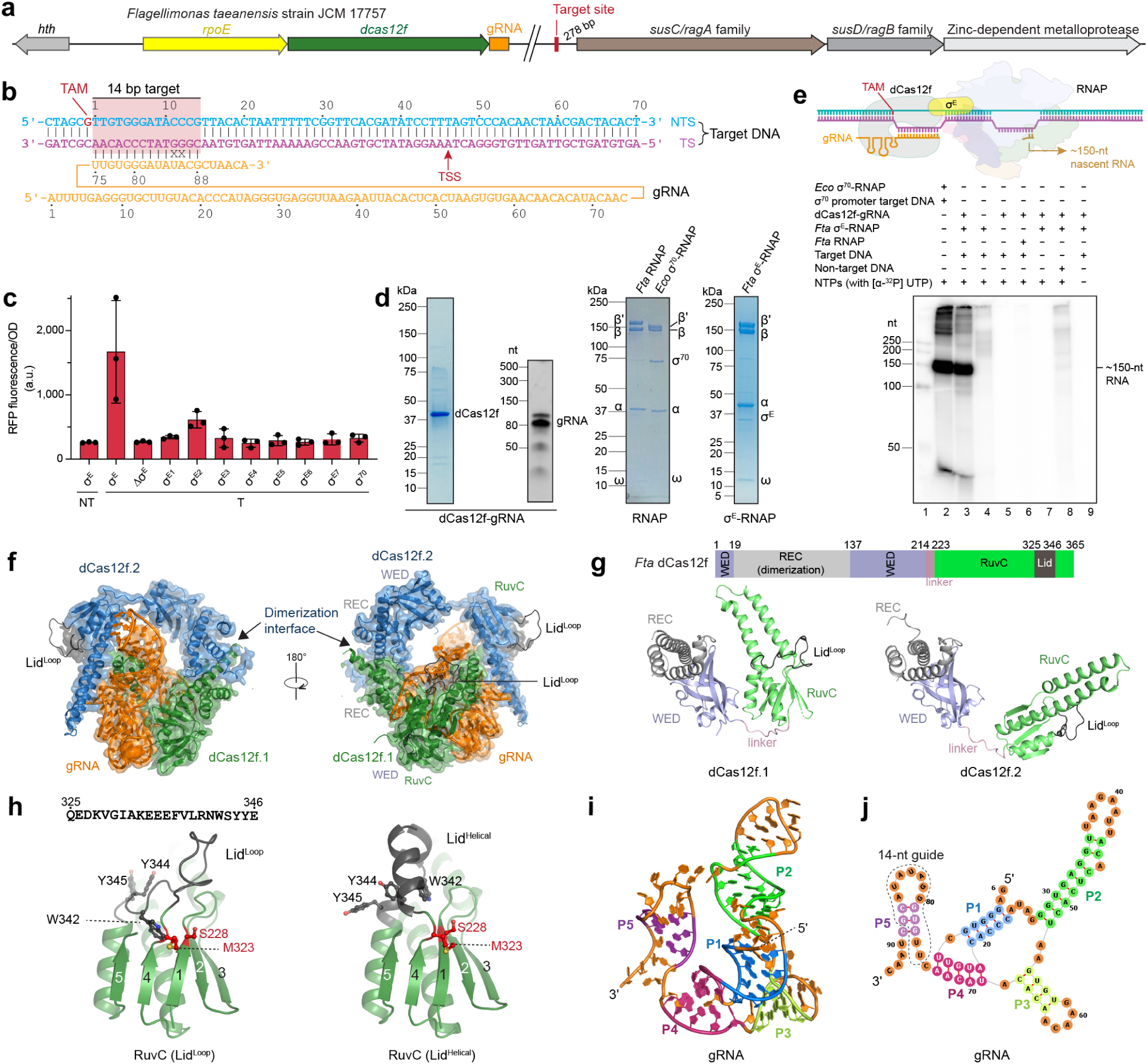
Biochemical characterization of the dCas12f-σ^E^ system and structure of dCas12f-gRNA. **a**, Genomic organi-zation of the dCas12f-σ^E^ operon in *F. taeanensis* (left), and the gRNA-matching DNA target site upstream of *susC/susD* mem-brane transport genes (right). **b**, Schematic showing base-pairing between the gRNA and target DNA, which includes native mismatches at positions 11–12. The downstream transcription start site (TSS) is indicated. **c**, OD-normalized RFP fluorescence from cellular transcriptional reporter assays using the indicated *Fta* σ^E^ homologs. dCas12f-gRNA complexes were presented in all conditions, and contained a targeting (T) or non-targeting (NT) guide. Data are shown as mean ± s.d. for *n* = 3 biologically independent samples. **d**, SDS-PAGE and Urea-PAGE analysis of purified dCas12f-gRNA, RNAP, and σ^E^-RNAP complex-es. Similar data were obtained for three independent preparations. **e**, *In vitro* transcription assays with radiolabeled NTPs using a 331-bp DNA template. The schematic indicates the TAM, TSS, and expected ∼150-nt RNA transcription product. RNA was only generated in the presence of dCas12f-gRNA, **σ**^E^, and RNAP on target DNA (lane 3); lane 2 shows an *E. coli* RNAP-σ^70^ control. Similar data were obtained from three biological replicates. **f**, Cryo-EM density map of the dCas12f-gRNA complex at 3.28 Å. **g**, Domain organization of dCas12f.1 and dCas12f.2, showing the REC, WED, RuvC, and lid motif domains. **h**, Struc-tures of the lid motif in ‘Loop’ (left) and ‘Helical’ (right) conformations. **i**, Cartoon representation of the experimental gRNA structure. **j**, Secondary structure diagram of the gRNA, highlighting key structural elements. Stem-loops are labeled P1–P5.

## RESULTS

### Biochemical basis of dCas12f-**σ**^E^ coupling

We focused on a *F. taeanensis* (*Fta*) strain that encodes one dCas12f-σ^E^ system alongside 26 additional σ^E^ homologs (*rpoE* gene) and 3 σ^70^ homologs (**Fig. 1a, Extended Data Fig. 1, and Supplementary Fig. 2**). Based on their phylogenetic distribution, we selected nine representatives for functional testing in *E. coli*, including the dCas12f-associated σ^E^, and developed a fluorescence-based reporter assay to monitor activity^6^, in which the native gRNA-matching target site is cloned upstream of RFP (**Fig. 1b**). The resulting data revealed that only the cognate, dCas12f-associated σ^E^ enabled RNA-guided transcription above background levels, whereas all other tested **σ** factors showed baseline activity comparable to the no-**σ** control (**Fig. 1c**). These data are consistent with a functional coupling that is exclusive to dCas12f and its genomically proximal *rpoE* gene product.

We next separately purified the *Fta* dCas12f-gRNA binary complex and the *Fta* RNAP core enzyme (α, β, β’, ω) using a heterologous *E. coli* expression system. Analysis by SDS-PAGE revealed an absence of contaminating *E. coli* RNAP components, suggesting that the purified products predominantly corresponded to homogeneous *Fta* RNAP (**Fig. 1d**). While *Fta* σ^E^ alone proved recalcitrant to purification, co-expression with RNAP yielded a stable σ^E^-RNAP complex (**Fig. 1d**). Similarly, while we were unable to isolate a homogenous complex from cells co-expressing σ^E^ with dCas12f-gRNA (**Extended Data Fig. 2a–c**), size-exclusion chromatography indicated that a stable dCas12f-σ^E^-RNAP supercomplex specifically formed in the presence of target DNA (**Extended Data Fig. 2d-i**). This latter result suggests that substrate DNA binding and R-loop formation triggers RNAP holoenzyme recruitment. *In vitro* transcription assays with radiolabeled NTPs confirmed dCas12f-dependent transcription and yielded an RNA product of ∼150 nucleotides (nt), consistent with a TSS located ∼46 base pairs (bp) downstream of the TAM **(Fig. 1e, Extended Data Fig. 2j, and Supplementary Table 1)**.

### Structure of dCas12f-gRNA

To elucidate the molecular basis of RNA-guided transcription, we first determined a 3.28 Å cryo-EM structure of the dCas12f-gRNA complex (**Fig. 1f, Extended Data Fig. 3, and Extended Data Table 1**). As with canonical Cas12f effectors such as *Un*Cas12f^19,20^ and *As*Cas12f^21,22^, *Fta* dCas12f assembles as an asymmetric dimer with a single gRNA (**Fig. 1f**).

dCas12f consists of an N-terminal region (1–214), further divided into the REC (recognition) and WED (wedge) domains, and a C-terminal RuvC domain (223–365) connected by a linker (**Fig. 1f,g**). The REC domain forms the dimerization interface, and consistent with previous nomenclature^19–22^, we designate the subunit that extensively interacts with the gRNA and binds the TAM as dCas12f.1, and the other as dCas12f.2. Domains within both subunits share identical folds, but the N- and C-terminal regions adopt different orientations due to linker variations. A key motif within the RuvC domain is the lid motif, a region between the β4 and β5 strands of the RuvC domain spanning residues 325–346 (**Fig. 1h**), which undergoes major conformational changes upon target DNA recognition, as discussed below.

The gRNA is naturally encoded downstream of *dcas12f* as a single-guide transcript^6^, in contrast to the natural dual-guide RNA comprising crRNA and tracrRNA used by canonical CRISPR-Cas12f nuclease systems^23^. The gRNA consists of a core structure with four based-paired regions (designated P1–P4) and a 14-nt 3’ guide region, which strikingly adopts an unexpected stem-loop structure P5 (**Fig. 1i,j, and Extended Data Fig. 3g,h**). This differs markedly from other Cas12 nucleases, where the seed sequence typically maintains a pre-ordered conformation to facilitate target DNA hybridization, with the rest of the guide region remaining flexible^20,24,25^. Conventional CRISPR-Cas systems rely on diversified guides that comprise unique, foreign DNA-derived spacers within the CRISPR array, whereas the dCas12f-σ^E^ system has evolved an inflexible guide region that targets a specific, conserved genomic site upstream of *susCD* for gene regulation^6^. This reduced variability may thus favor the formation of the P5 stem-loop structure, which contains base pairs between residues 77–79 and 87–89 (**Fig. 1i,j**). This region must unwind to enable RNA-DNA heteroduplex formation, likely imposing an energetic barrier that enhances target specificity.

### Structure of DNA-bound dCas12f-gRNA

To investigate the mechanism of RNA-guided DNA binding, we next investigated the ternary com-plex comprising dCas12f, gRNA, and target DNA (**Fig. 2a,b, Extended Data Fig. 4, and Extended Data Table 1**). Overall, the ternary complex resembles *As*Cas12f-gRNA-DNA and *Un*Cas12f-gRNA-DNA complexes^19–21^, but with minimized protein and gRNA constituents (**Supplementary Fig. 3**). The TAM duplex binds within a groove formed by the REC and WED domains of dCas12f.1, which make extensive contacts with the phosphate backbone of the non-target strand (**Fig. 2c,d**), and alanine substitutions of key residues involved in this interaction (Y89 and R90) reduce RNA-guided transcription by roughly 50% (**Fig. 2e, and Supplementary Figs. 4 and 5**). Strand separation is facilitated by H110 and M85, which stack against a G-C pair at position −1, and recognition of this minimal 5’-G TAM is mediated by a hydrogen bond between the S106 main-chain carbonyl group and the C(−1) base of the target strand (**Fig. 2f**). Additionally, Q144 forms a hydrogen bond with A(−4) of the non-target strand **(Fig. 2c,d)**. While the TAM binding site in Cas12f.2 is accessible, no DNA is bound to this site, consistent with the paradigm in which TAM (or PAM) interactions are transient and only stabilized upon RNA-DNA hybrid formation^26^ (**Fig. 2a,b**).

**Figure 2.**
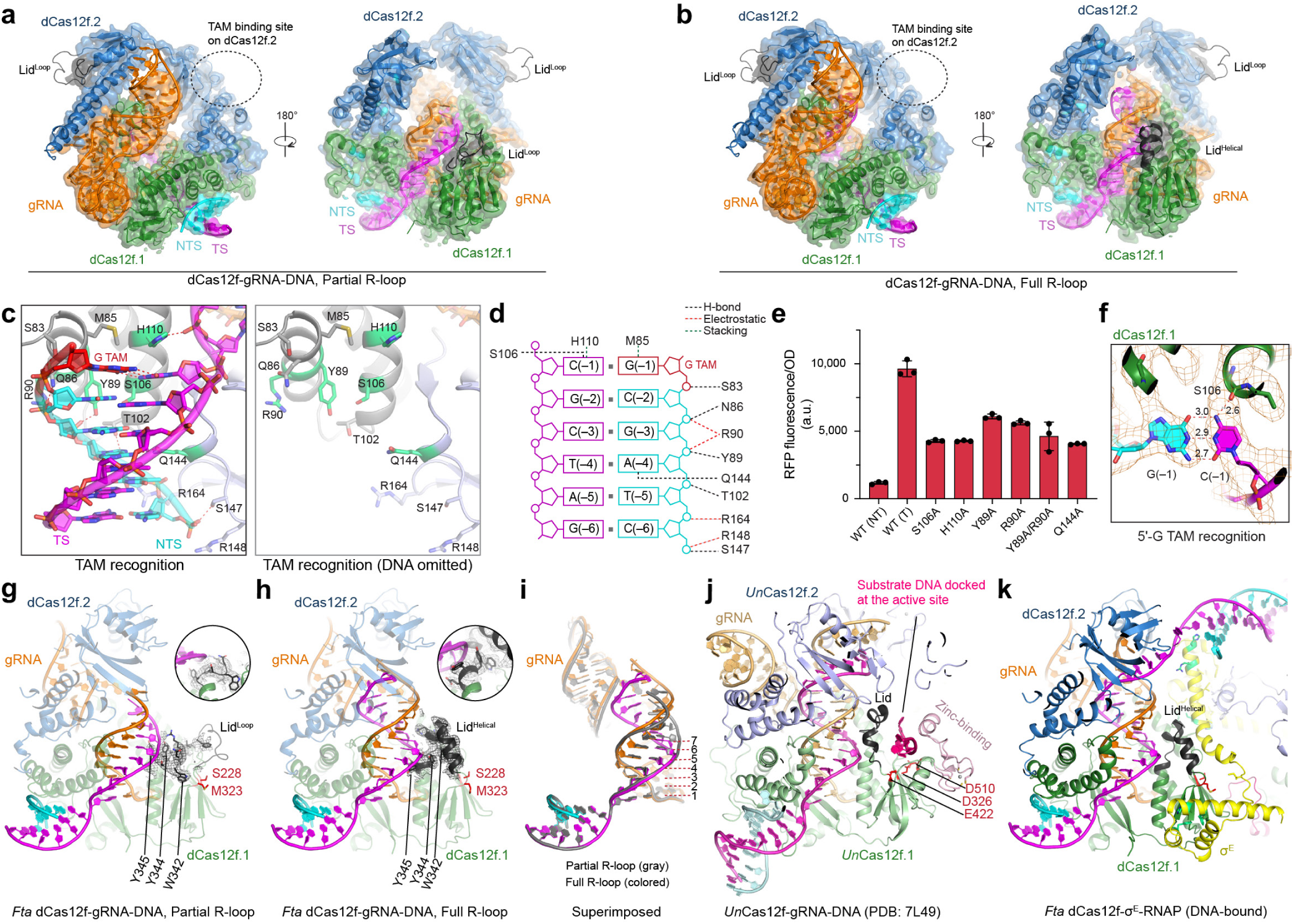
Structure of the dCas12f-gRNA-target DNA complex. **a**, Cryo-EM density map of dCas12f-gRNA-DNA in a partial R-loop state. **b**, Cryo-EM density map of dCas12f-gRNA-DNA in a full R-loop state. **c**, Interaction between the TAM duplex and dCas12f.1. **d**, Schematic of interactions between the TAM duplex and dCas12f.1. **e**, OD-normalized RFP fluorescence from cellular transcriptional reporter assays using the indicated dCas12f mutants that perturb TAM duplex recognition. Data are shown as mean ± s.d. for *n* = 3 biologically independent samples. **f**, Recognition of the 5’-G TAM, highlighting hydrogen bonding between the base of C(−1) and the backbone carbonyl oxygen of S106. **g**, Lid motif conformation in the partial R-loop state. **h**, Lid motif conformation in the full R-loop state. Insets in **g** and **h** show focused views of key residues in the lid motif undergoing conformational rearrangement, along with the corresponding cryo-EM density. **i**, Structural superimposition of RNA-DNA duplex in partial and full R-loop states. **j**, Lid motif conformation in *Un*Cas12f-gRNA-DNA complex for com-parison. **k**, Lid motif conformation in the dCas12f-gRNA-DNA-σ^E^-RNAP complex.

**Figure 3.**
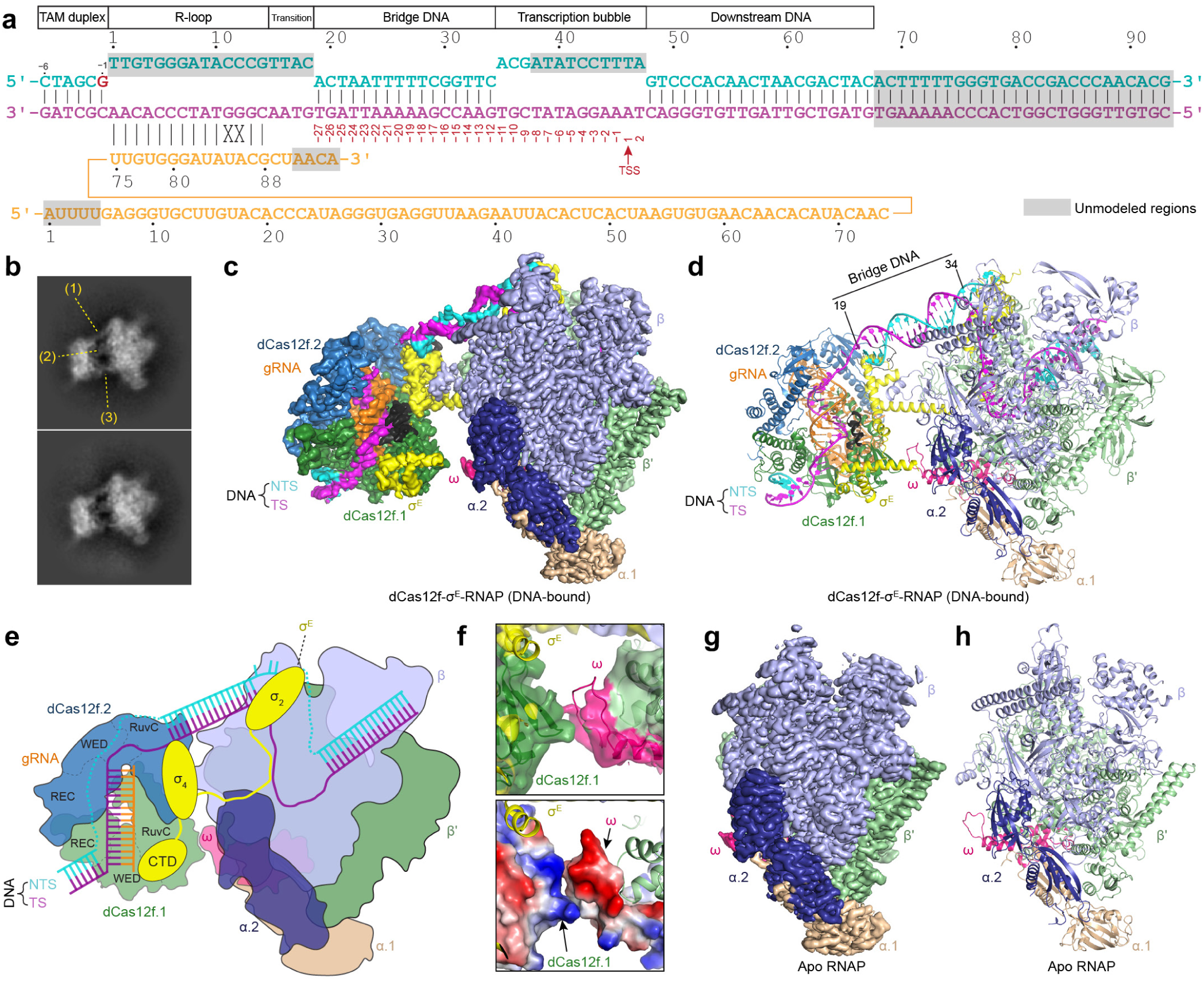
Structures of the DNA-bound dCas12f-σ^E^-RNAP complex and apo RNAP. **a**, Schematic showing the target DNA and gRNA in the DNA-bound dCas12f-σ^E^-RNAP structure. The target DNA features two DNA bubbles, one mediated by the dCas12f-gRNA R-loop and the other mediated by the RNAP transcription bubble; the TSS is located at position 46 of the template strand. **b**, Two representative 2D class averages of cryo-EM images of the DNA-bound dCas12f-σ^E^-RNAP complex. **c**, Cryo-EM density map of the DNA-bound dCas12f-σ^E^-RNAP complex. **d**, Cartoon representation of the atomic model. **e**, Schematic of the DNA-bound dCas12f-σ^E^-RNAP structure. **f**, Contact between dCas12f.1 and the RNAP ω subunit, showing the cryo-EM map displayed as a surface representation (top) and the surface potential of dCas12f.1 and the ω subunit (bottom). **g**, Cryo-EM map of apo RNAP. **h**, Atomic model of apo RNAP.

**Figure 4.**
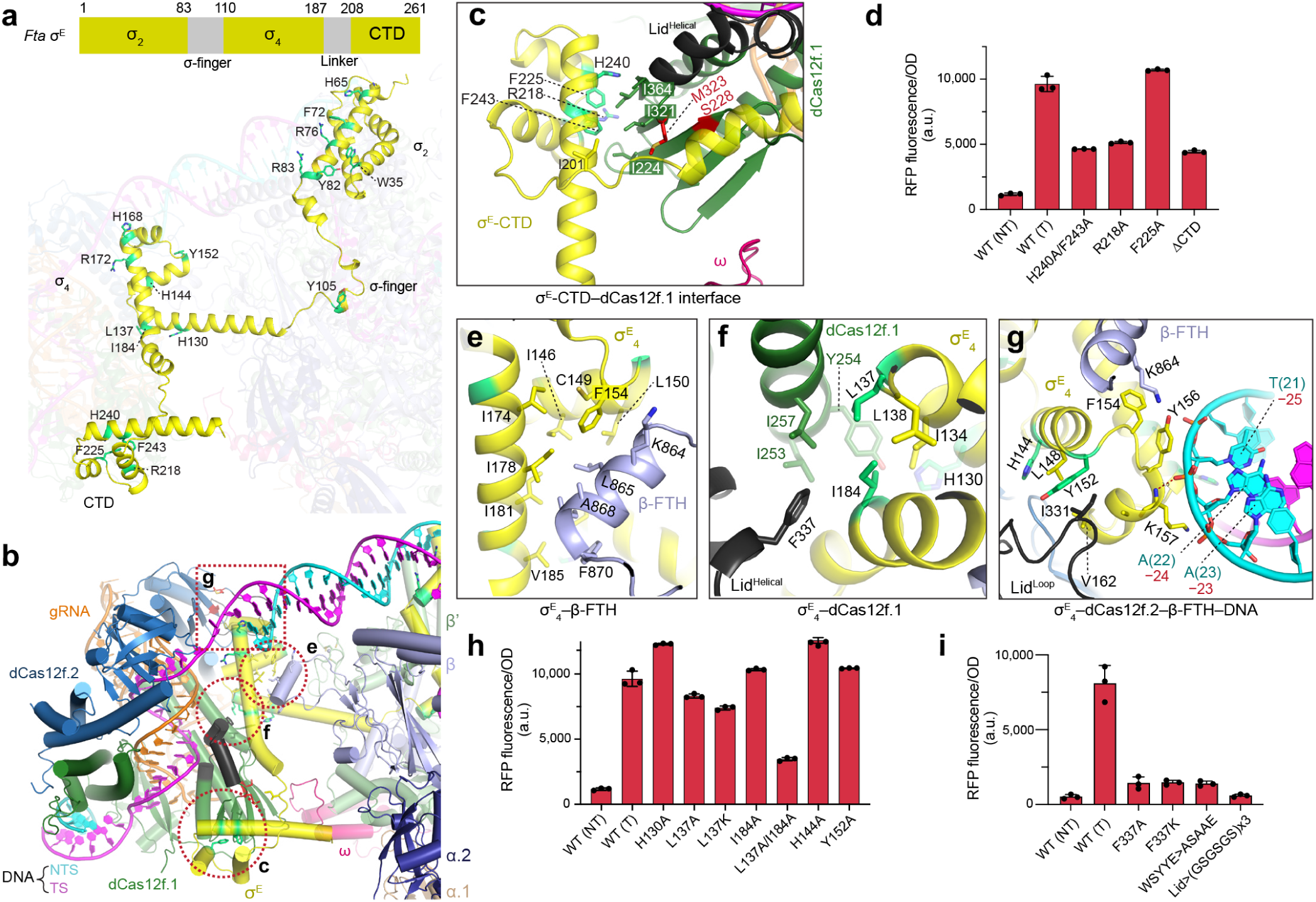
Interactions between dCas12f and σ^E^. **a**, Structure of σ^E^ in cartoon representation, with the remainder of the DNA-bound dCas12f-RNAP complex shown in transparency, for clarity. The σ^E^ domain organization is shown at the top. **b**, Overall view showing the interactions between σ^E^ and other components, with magnified inset regions indicated with dotted lines. **c**, Interactions between the σ^E^ CTD and dCas12f.1. **d**, OD-normalized RFP fluorescence from cellular transcriptional reporter assays using the indicated σ^E^ CTD mutants. **e**, Interactions between the σ^E^ domain and β-flip-tip-helix (β-FTH). **f**, Interactions between the σ^E^ domain and dCas12f.1. **g**, The σ^E^ domain is encircled by β-FTH, the lid motif of dCas12f.2, and DNA. **h**, OD-normalized RFP fluorescence from cellular transcriptional reporter assays using the indicated σ^E^ mutants. **i**, OD-normalized RFP fluorescence from cellular transcriptional reporter assays using the indicated dCas12f lid-motif mutants. Data in **d, h**, and **i** are shown as mean ± s.d. for *n* = 3 biologically independent samples. Residues highlighted in green (panels a, c, e, f, and g) indicate positions tested by mutagenesis.

**Figure 5.**
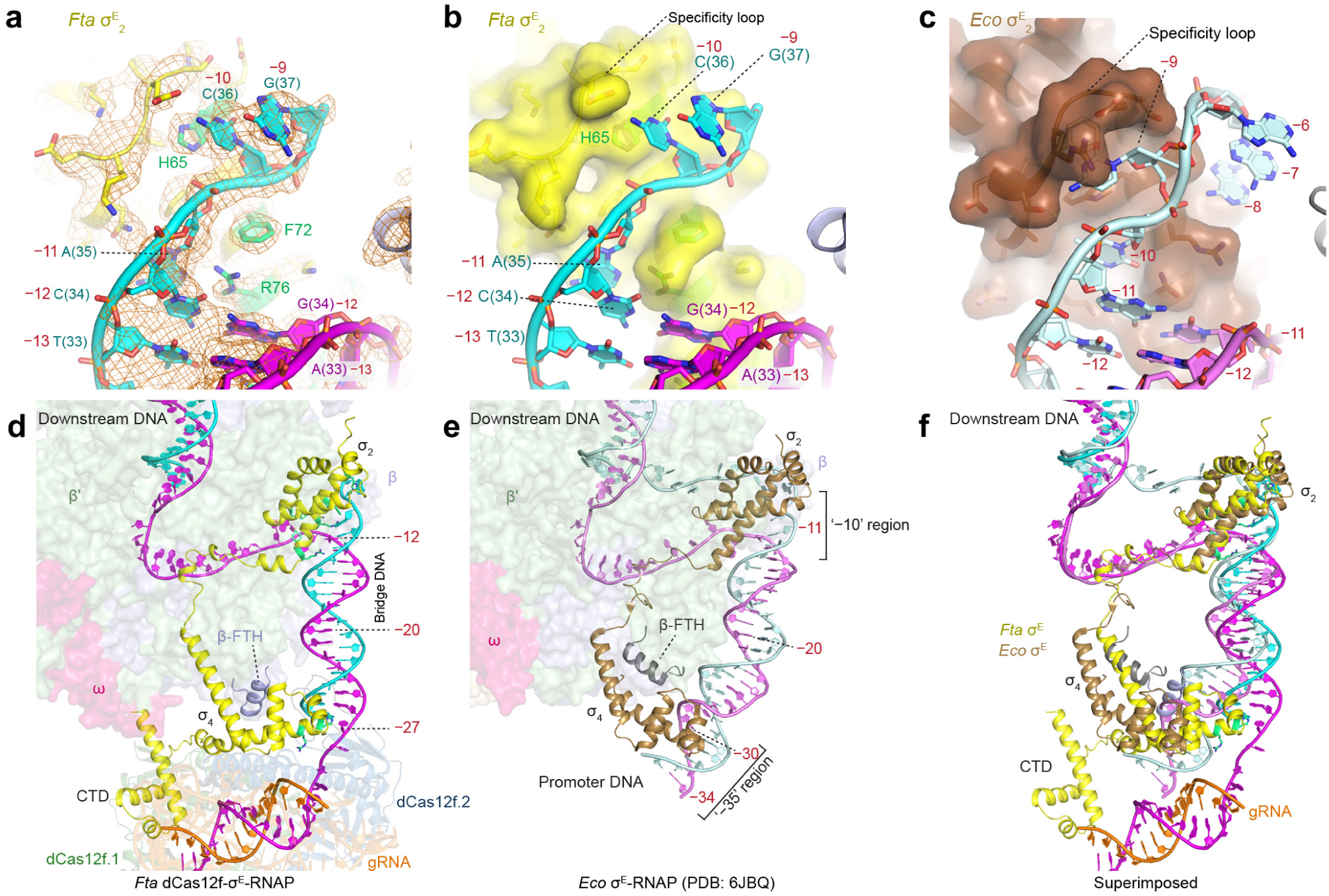
Target DNA interactions and comparison with canonical σ^E^ factors. **a**, Interactions between *Fta* σ^E^ and the −10 region of DNA; the cryo-EM map is shown in mesh. **b**, Interactions between *Fta* σ^E^ and the −10 region of DNA, with the σ^E^ surface displayed. **c**, Interactions between the *Eco* σ^E^ and the −10 region of DNA, shown as in **b**, highlighting the pocket formed by the specificity loop. **d**, Interactions between *Fta* σ^E^ and the target DNA. **e**, Interactions between *Eco* σ^E^ and the target DNA. **f**, Superimposed structure comparing dCas12f-associated *Fta* σ^E^ (yellow) and non-dCas12f-associated *Eco* σ^E^ (brown).

Hybridization of the guide RNA and target DNA occurs subsequent to TAM recognition, and 3D classification of the ternary complex revealed two distinct conformational states. The majority of particles (∼88%) adopt a partial R-loop conformation, with only a minor subset (∼12%) favoring a complete R-loop (**Fig. 2g,h, Extended Data Fig. 4f**). The partial R-loop structure features a 7-bp RNA-DNA heteroduplex, and although RNA and DNA densities at positions 8 and 9 are observed, they deviate from A-form duplex geometry and fail to form base pairs (**Fig. 2g,i, Extended Data Fig. 4g–j**). In the full R-loop structure, base pairs at positions 7–9 reorganize to adopt canonical A-form structure and the duplex extends to position 14 (**Extended Data Fig. 4h,j**), corroborating accompanying *in vivo* data from RNA immunoprecip-itation sequencing (RIP-seq) and transcriptional activity assays^6^ that showed that efficient transcription requires guide lengths greater than 10 nt. A twist in the target DNA forms a wobble base pair at position 10, potentially accommodating mismatches at positions 11 and 12 (**Extended Data Fig. 4j**), and densities for DNA and RNA are undetectable beyond position 14. *In vitro* transcription assays showed that restoring complementarity at positions 11–12 modestly increased activity (**Extended Data Fig. 2j**), indicating that these mismatches do not substantially alter the biological outcome of the system.

The transition between the partial and full R-loop states involves coordinated conformational changes in the RuvC domain, particularly in the lid motif. In the partial R-loop state, where we refer to the motif as lid^Loop^, only the C-terminal portion of the lid motif (residues 337–346) adopts a defined structure, while the N-terminal region (residues 325–336) is disordered (**Fig. 2g, and Supplementary Fig. 4**). Two consecutive tyrosines (Y344 and Y345) are positioned beneath the target DNA strand and sterically block position 8 from hybridizing with the gRNA in an A-form configuration. Upon RNA-DNA heteroduplex extension beyond position 7 in the full R-loop state, both tyrosines rearrange into an extended α-helical structure formed by residues 347–358. In the full R-loop state, the entire lid motif (referred to as lid^Helical^) becomes ordered, forming stable helical structures that interact with the target DNA **(Fig. 2h,i)**. These structures therefore reveal a likely mechanism whereby the lid motif functions as a sensor for R-loop formation. In canonical Cas12f nucleases, the lid motif guides the DNA substrate into the RuvC catalytic center, positioning it for cleavage^20^ (**Fig. 2j**), whereas for dCas12f-σ^E^ systems, we hypothesized that it would instead play a critical role in σ^E^ recruitment (**Fig. 2k**).

### Structure of dCas12f-σ^E^-RNAP holoenzyme

To directly resolve the mechanism of σ^E^ and RNAP recruitment during transcription initiation, we next determined the structure of the fully assembled dCas12f-σ^E^-RNAP nucleoprotein complex comprising dCas12f-gRNA, a 99-bp target DNA, σ^E^, and *Fta* RNAP. The target DNA corresponds to the native site located ∼278 bp upstream of the *sucC* gene and comprises a 6-bp upstream region containing the TAM, a 14-bp target site matching the gRNA, and 79 bp of downstream sequence. The dCas12f-σ^E^-RNAP complex was refined to an average resolution of 3.30 Å from 11,161 particles, and 3D classification also allowed us to determine the apo RNAP structure to 2.95 Å from 171,804 particles (**Extended Data Fig. 5**, **Extended Data Table 1**). Optimized DNA constructs incorporating mismatches to favor DNA bubble formation within the RNAP core improved the overall resolution to 2.88 Å (**Extended Data Fig. 6, Extended Data Table 1**), and focused refinement using a mask around the dCas12f-gRNA region yielded a local resolution of 3.34 Å (**Extended Data Fig. 6e**). The final maps from matched and mismatched dsDNA were similar, and we used the higher-resolution mismatched dsDNA map for model interpretation. Overall, the final map showed excellent density, allowing model building for nearly all protein subunits of the complex (**Extended Data Fig. 7**). However, the α subunit C-terminal domains, known to interact with the UP element of canonical promoter DNA^27^, are not resolved in the map, indicating their inherent flexibility. The lineage-specific insertions in the β and β’ subunits, which likely interact with transcriptional factors^28^, are visible but at lower resolution (**Extended Data Fig. 7a,b**).

The DNA substrate can be divided into six regions: the TAM duplex, R-loop, transition region, bridge DNA, transcription bubble, and downstream DNA **(Fig. 3a)**. The 6-bp TAM duplex and 14-bp RNA-DNA heteroduplex within the R-loop are well-resolved, although density for the non-target DNA strand is largely absent (**Extended Data Fig. 7j, 8a,b**). The bridge DNA, which spans between dCas12f and the RNAP core, adopts a B-form conformation with well-resolved major and minor grooves (**Fig. 3b–d**), but weak density between the R-loop and bridge DNA suggests structural flexibility in this transition region, preventing precise sequence tracing (**Extended Data Fig. 7j**). Within the transcription bubble itself, we modeled a 13-nt sequence for the template strand, whereas only the first 3 nt are visible for the non-template strand (**Extended Data Fig. 7j**); this observation aligns with the typical transcription bubble length of 12–14 bp observed in homologous bacterial RNAP structures^29–33^. Finally, the downstream DNA is modeled as a 17-bp B-form duplex.

To provide a mechanistic framework for transcription initiation, we modeled a 4-nt transition region (positions 15–18) based on structural constraints and cryo-EM densities from the unsharpened map (**Extended Data Fig. 7j**). This allowed confident tracing of the template DNA strand sequence up to position 67 in the downstream DNA, yielding a model that aligns well with the observed density and is biologically plausible, as it places position 46, the most frequently observed TSS from RNA-sequencing data^6^, within the RNAP catalytic center. We note, however, that minor shifts in conformation may occur given the inherent flexibility of DNA^34^, and this could readily explain the observation of context-dependent TSSs located 45–48 bp downstream of the TAM site^6^. Collectively, these findings suggest that our structure represents a biologically relevant state of RNA-guided transcription driven by the dCas12f-σ^E^ system.

The final structural model reveals a pronounced bipartite architecture, consisting of a well-ordered RNAP core region bound to the σ_2_ domain of σ^E^, and a smaller dCas12f-gRNA region that engages with the σ_4_ and C-terminal domains (CTD) of σ^E^ (**Fig. 3a–e**). These two regions are physically connected by three distinct structural elements: the 16-bp bridge DNA (**Fig. 3b–d**), the elongated σ^E^ α-helix that links its RNAP-bound σ_2_ domain to its dCas12f-bound σ_4_ domain (**Fig. 3b–d**), and electrostatic interactions between the dCas12f.1 RuvC domain and the RNAP ω subunit (**Fig. 3f**).

### *Fta* RNAP ω subunit contacts dCas12f

RNAP and σ factors canonically function together for faithful promoter recognition^8^, but in dCas12f-σ^E^ systems, RNAP also promotes key interactions between DNA-bound dCas12f and σ^E^. Overall, the core subunits closely resemble other bacterial RNAP structures, such as those from *E. coli* and *M. tuberculosis*^29,33^ (**Extended Data Fig. 9a–d**), with the characteristic ‘crab-claw’ shape formed by the large β and β’ subunits (**Extended Data Fig. 10**). These two subunits form a structural platform that is stabilized by peripheral α subunits (α1 and α2) and the ω subunit, while also coordinating σ^E^ recruitment. All key structural elements in the β and β′ subunits that are critical for transcription are conserved in *Fta* RNAP, as observed in other bacterial RNAP structures, including the clamp domain that accommodates the target DNA and engages σ^E^, the NTP entry funnel that channels incoming substrates, the active-site cleft responsible for RNA synthesis, and the positively charged bridge helix (BH) that plays a crucial role in catalysis and RNA-DNA translocation^33,35^ **(Extended Data Fig. 10)**. Notably, the β flat tip helix (β-FTH) plays a crucial role in engaging the σ_4_ domain of σ^E^ (ref. ^36^), as detailed below.

As the smallest RNAP core subunit, ω primarily interacts with the β’ subunit for proper β’ subunit folding^37^ and have chaperone-like role in RNAP assembly^38^. Compared to *E. coli* ω, *Fta* ω contains a negatively charged insertion (residues 48–82) that adopts a helix-loop-helix structure. This insertion appears unique among Bacteroidota (**Extended Data Fig. 9e**), is conserved in genomes containing dCas12f-σ^E^ systems (**Extended Data Fig. 9f,g, and Supplementary Fig. 6**), and engages the basic RuvC domain of Cas12f.1 (**Fig. 3f**). A DALI search identified a similar ω subunit insertion in *Porphyromonas gingivalis* RNAP (PDB: 8DKC)^39^ (**Extended Data Fig. 9g**), but dCas12f is notably absent from the genome. *P. gingivalis*, like *F. taeanensis*, belongs to the bacterial phylum *Bacteroidetes*, thus suggesting that the conserved structural insertion reflects a shared phylogenetic origin and was co-opted for dCas12f interactions.

### The σ^E^–dCas12f interactions

σ^E^ is recruited to target-bound dCas12f via both protein–protein and protein-DNA interactions (**Fig. 4a,b**) that provide critical insights into RNA-guided transcription. The σ^E^ CTD, which is unique to dCas12f-associated σ^E^ and absent in other σ^E^ paralogs^6^, binds directly to the RuvC domain of dCas12f.1, complementing its partially truncated structure (**Fig. 4c**). The interface centers on a hydrophobic core formed by σ^E^ residues F225 and F243, and hydrophobic residues in the dCas12f.1 RuvC domain. The interaction is further strengthened by hydrogen bonding interactions, including those mediated by H240 and R218 (**Fig. 4c, and Supplementary Fig. 7**). Deletion of residues 201–265 in the σ^E^ CTD, or alanine substitutions in this region (R218A and H240A/F243A), led to a ∼50% reduction in transcriptional activity compared to wild type in cellular reporter assays (**Fig. 4d, and Supplementary Fig. 5**). Consistently, biochemical reconstitution assays led to a pronounced loss of activity for these variants, despite comparable assembly with RNAP (**Extended Data Fig. 2k,i**).

The σ_4_ domain acts as a linchpin, contacting dCas12f, the target DNA, and the β-FTH **(Fig. 4b,e,f,g)**. Two hydrophobic residues, I184 and L137, form key contacts with the dCas12f.1 lid^Helical^ motif (**Fig. 4f**). Substitution of L137 with alanine or lysine significantly reduced transcription activity in cellular assays, and the L137A/I184A double mutation substantially reduced transcriptional activity (**Fig. 4h**). Moreover, mutations of key residues within the lid motif of dCas12f (e.g., F337A, F337K), or replacement of the entire lid motif with a flexible GSGSGS linker, severely impaired transcription activity in both cellular reporter assays and *in vitro* transcription assays **(Fig. 4i and Extended Data Fig. 2m,n)**. In addition to this key contact, the σ_4_ domain engages with the target DNA and dCas12f.2 simultaneously via residues H144 and Y152 of the latter’s lid motif (**Fig. 4g**). The WT-level activity of H144A and Y152A mutants, however, suggests that this interaction is ancillary rather than critical. Notably, the σ_4_ domain engages with the target DNA through minor groove contacts (**Fig. 4b**, **Extended Data Fig. 8f**), contrasting with its canonical role in recognizing −35 promoter elements via major groove interactions in typical σ factors^29,40^. One exception is the σ^I^ (SigI) factor, which recognizes the −35 region through both the major groove and A-tract minor groove, likely enhancing binding stability via increased interface area^41^. Minor groove recognition follows a similar pattern, with a positively charged histidine inserting into the groove and positively charged residues interacting with the phosphate backbone, commonly found in protein-DNA interactions^42^. This altered DNA binding mode suggests that σ_4_ primarily stabilizes the complex rather than conferring sequence specificity, allowing dCas12f-gRNA to provide target recognition function.

### σ^E^-DNA interactions

The σ_2_ domain plays key roles in transcription initiation, particularly in DNA unwinding and strand separation at the −10 region^9^. In our structure, R76 packs against the last base-paired position (G-C pair at position −12 from the TSS), facilitating strand separation (**Fig. 5a,b, Extended Data Fig. 8c,d**). Upon DNA melting, both strands are stabilized by distinct protein interactions to prevent re-annealing: the non-template strand is primarily stabilized by the σ_2_ domain (**Fig. 5a,b**), while the template strand is stabilized through interactions with both σ^E^ and the RNAP β-protrusion (**Extended Data Fig. 8c,d**). Overall, multiple key residues contribute to maintaining a melted DNA state, though single mutations did not reduce transcriptional activity, suggesting that individual perturbations are tolerated under these experimental conditions (**Extended Data Fig. 8g**).

In primary σ factors, promoter sequence specificity is largely determined by the −10 element (Pribnow box ; consensus TATAAT from positions −12 to −7) on the non-template strand, with the −11A and −7T nucleotides being the critical determinants^12,29–31,43,44^. By contrast, in *E. coli* σ^E^, a specificity loop between two helices forms a binding pocket that recognizes the flipped-out nucleotide^29^ (**Fig. 5c**). In our structure, however, no obvious binding pocket is present for recognizing a flipped nucleotide, and H65 instead packs against C(−10) and G(−9) (**Fig. 5b**), likely stabilizing the melted bases and preventing re-annealing without sequence specificity. Consistently, perturbation of the −10 region (or the −7 region of Flavobacteria within the phylum *Bacteroidetes* ^45,46^) had minimal effect on RNA-guided transcription^6^, and the H65A substitution caused a moderate reduction in activity **(Extended Data Fig. 8g)**. These observations support a model in which H65 serves as an important contributor within an interaction network that stabilizes the melted transcription bubble in a sequence-independent manner.

The σ finger extends into the catalytic cleft, functioning similarly to the primary σ_3.2_ domain (σ_3/4_ linker) of σ^70^ (ref. ^12,47,48^) by pre-organizing the template strand for subsequent *de novo* RNA synthesis, as is also observed in homologous ECF sigma factors^29–31^. In our structure, Y105 from the linker packs against G(−4) of the template strand (**Extended Data Fig. 8e**).

In comparing our *Fta* dCas12f-σ^E^-RNAP structure with the *E. coli* σ^E^-RNAP complex, notable differences emerge. Whereas the σ_4_ domain typically recognizes the −35 region through major groove contacts^29,40^, in our structure it instead engages with the minor groove (**Fig. 5d–f**). Additionally, the bridge DNA (corresponding to the upstream DNA in the *E. coli* σ^E^-RNAP complex) adopts a distinct angle by rotating away from the RNAP β’-ZnF (**Fig. 5d–f, Extended Data Fig. 9a-c**), thus deviating from the typical configuration observed in standard transcription initiation complexes^32,47,49,50^. The bridge DNA region is notably A-T rich, with a continuous A-T sequence spanning positions 21–28. The A-T rich bridge may act as a flexible hinge, allowing the DNA to adopt a conformation favorable for interaction with the dCas12f-σ^E^-RNAP complex.

## DISCUSSION

Altogether, our results reveal an unprecedented mechanism of transcription initiation in dCas12f-σ^E^ systems, in which protein-DNA promoter recognition is replaced by RNA-DNA complementarity, enabling a flexible and programmable mode of defining transcription start sites (TSSs). The dCas12f-gRNA complex binds upstream of the TSS at a position analogous to the −35 promoter region in canonical bacterial transcription^30,49,50^ (or the −33 region of Flavobacteria within the phylum *Bacteroidetes*^45,46^), leading to subsequent recruitment of the σ^E^-RNAP complex. The σ_4_ domain, which is usually responsible for −35 sequence recognition, plays a distinct role by serving as a structural adaptor that bridges dCas12f and RNAP through critical engagement of the minor groove of the bridge DNA. Meanwhile, the σ_2_ domain retains its conventional role in core RNAP interactions and DNA melting at the −10 region, though the sequence specificity enforced is substantially reduced. Together, these structural adaptations exemplify striking co-evolution between RNA-guided, nuclease-dead Cas12f proteins and extracytoplasmic function sigma factors, that enabled a novel mode of transcriptional regulation.

Our structural analysis identified three key interaction points that establish the dCas12f-σ^E^-RNAP complex: a bridge DNA-mediated connection, direct interactions between dCas12f and σ^E^_4_ that are further stabilized through the minor groove DNA contacts, and direct dCas12f-RNAP core interactions through the ω subunit. This latter interaction may represent a key feature to accommodate RNA-guided transcription. Interestingly, the ω subunit has also been exploited in engineered CRISPRa systems, where fusion of ω to dCas9 or TALE recruits RNAP to activate transcription in the presence of promoters^3,51^. The natural dCas12f-σ^E^ system achieves a similar outcome without covalent fusions, relying instead on specific protein-protein interactions.

The σ^E^ C-terminal domain (CTD) extension is a unique feature of dCas12f-associated systems that distinguishes it from canonical σ factors (**Extended Data Fig. 1**). It binds the truncated, nuclease-inactive RuvC domain of dCas12f, likely enhancing the affinity between the RNA-guided effector and the σ^E^-RNAP complex. Given its positioning, we speculate that this critical σ^E^-Cas12f.1 interaction functions as an initial docking mechanism, facilitating the recruitment of σ^E^-RNAP to the bound target DNA. This protein-protein interaction may provide a more efficient means of transcriptional activation compared to conventional sigma factor-mediated promoter search mechanisms^52^. Our data also highlight the conformational coupling between R-loop formation and transcription complex assembly. Specifically, conformational changes of the dCas12f lid motif function as a molecular switch that ensures transcription initiation occurs exclusively at targeted sites, likely enhances both specificity and regulatory control in cellular contexts.

In conclusion, our work establishes a structural framework for RNA-guided transcription, elucidating key interactions between an RNA-guided protein and the σ^E^-RNAP holoenzyme complex. These insights provide new opportunities for developing programmable transcriptional regulators for precise gene expression control.

## METHODS

### Molecular constructs

The initial plasmid construct (pSL6726) that encodes *F. taeanensis* dCas12f, gRNA, and σ^E^ (*rpoE* gene) was synthesized by GenScript^6^. *dcas12f* and *rpoE* genes were individually subcloned into the bacterial expression vector pET28-MKH8SUMO (Addgene #79526). In addition, both genes were cloned together into a single pET28-MKH8SUMO construct **(Extended Data Fig. 2a)**. For co-expression with dCas12f or dCas12f-σ^E^, the gRNA with T7 promoter and T7 terminator was subcloned into a pUC19 vector. However, expression of σ^E^ alone or together with dCas12f and gRNA did not produce soluble σ^E^ **(Extended Data Fig. 2b)**. For co-expression with the His-tagged *Fta* RNAP, the His -tag in the *rpoE* vector was subsequently removed. Genes encoding *Fta* RNAP subunits were also synthesized by GenScript and subsequently cloned into the pACYCDuet vector (*rpoA*_*rpoB* in MCS-1 with a His_6_-tag on *rpoA*; *rpoC*_*rpoZ* in MCS-2).

### Mutagenesis

Single amino acid mutations were introduced using the QuikChange site-directed mutagenesis method. Multiple amino acid mutations were generated by ligating inverse PCR-amplified backbone with mutation-bearing DNA oligonucleotides using the In-Fusion Cloning Kit (Clontech). All mutations were verified by Sanger sequencing.

### Protein expression and purification

dCas12f and gRNA, or the dCas12f-RpoE construct and gRNA, were co-expressed in Rosetta(DE3)pLysS cells (Novagen #70956). Notably, although pET28-MKH8SUMO (ColE1-type origin) and pUC19 (modified ColE1/pMB1 origin) belong to the same incompatibility group, both plasmids were maintained under short-term dual antibiotic selection^53^, allowing successful co-expression. σ^E^ and RNAP were co-expressed in BL21-AI cells (Thermofisher # C607003).

Cells were cultured in Terrific Broth (TB) to an OD_600_ of 0.6. Protein expression was induced with 0.3 mM IPTG, and for BL21-AI cells, 0.2% arabinose was added. Cultures were incubated overnight at 16 °C. Cells were harvested and resuspended in lysis buffer (25 mM Tris-HCl pH 7.6, 500 mM NaCl, 5% glycerol) supplemented with 1 mM PMSF and 5 mM β-mercaptoethanol, then lysed by sonication. The lysate was clarified by centrifugation, the pellet was solubilized in 8 M urea for inclusion body detection, and the supernatant was applied to Ni-NTA resin. After extensive washing with lysis buffer containing 30 mM imidazole, target proteins were eluted with lysis buffer containing 250 mM imidazole.

The His-SUMO tag from dCas12f and the SUMO tag from σ^E^ were removed by overnight digestion with SUMO protease at 4 °C. The dCas12f-gRNA eluate was diluted in buffer containing 25 mM Tris-HCl pH 7.6, 125 mM NaCl, and 5% glycerol, then loaded onto a Heparin column (Cytiva) and eluted using a linear NaCl gradient (0.1 to 1 M). The σ^E^-RNAP eluate was diluted in the same buffer but purified using an HP Q column (Cytiva). Following concentration, proteins were further purified by size exclusion chromatography (SEC) using a Superose 6 Increase 10/300 GL column (Cytiva) in equilibration buffer containing 25 mM Tris-HCl pH 7.6, 250 mM NaCl, 5% glycerol, and 2 mM DTT. Purified fractions were concentrated and stored at -80 °C.

The *E. coli* RNAP–σ^70^ holoenzyme (NEB, M0551S) was purchased as a control for detecting chimeric *Fta*-*Eco* RNAP formation.

### *In vitro* transcription assay

The DNA templates for *in vitro* transcription were PCR-amplified from synthesized gBlocks (IDT) (sequence details in **Supplementary Table 1**) and cleaned by QIAquick PCR Purification Kit (Qiagen). The template used in **Figure. 1e** and used for testing all mutants *in vitro* is an optimized DNA containing an AT-rich region between TAM and TSS (**Supplementary Table 1)**. The WT template DNA used in **Extended Data Fig. 2j** is derived from the PAM-TSS region in the native target locus. The template DNA (400 ng) was incubated with dCas12f-gRNA (1.2 μM) and σ^E^-RNAP (0.6 μM) on ice in the reaction buffer (20 mM Tris pH 7.6, 100 mM KCl, 5 mM MgCl_2_, 10 mM DTT, 0.01% Triton® X-100) for 15 minutes. Transcription was initiated by adding radiolabeled nucleotide triphosphates (NTPs) (final concentrations: 250 μM each of CTP, GTP, and ATP; 25 μM UTP; and 0.5 μCi [α-³²P] UTP [3000 Ci/mmol, 10 mCi/mL]). Reactions proceeded for 45 minutes at 37 °C and were terminated by adding 20 μL 2X loading buffer (8 M urea, 50 mM EDTA, 0.02% xylene cyanol, 0.02% bromophenol blue) followed by incubation at 95 °C for 10 minutes. RNA transcripts were resolved by 15% Urea-PAGE (Bio-Rad) and analyzed by storage-phosphor scanning (Bio-Rad).

The RNA marker was prepared in-house by transcribing five template DNAs carrying a T7 promoter **(Supplementary Table 1)** that yielded 250-nt, 200-nt, 150-nt, 100-nt, and 50-nt RNA products, using the HiScribe® T7 High Yield RNA Synthesis Kit (NEB # E2040S) and radiolabeled NTPs as described above. The *in vitro* transcription of control ecoRNAP-σ^70^ was performed similarly as described above, with slight modification on the reaction buffer (40 mM Tris-HCl pH 7.6, 150 mM KCl, 10 mM MgCl_2_, 1 mM DTT, 0.01% Triton® X-100) and using a σ^70^-specific template DNA **(Supplementary Table 1)**.

### Complex assembly

The dCas12f-gRNA-target DNA ternary complex was reconstituted by incubating dCas12f-gRNA with a 60-bp target DNA (**Supplementary Table 1**) at a 1:1.3 ratio for 30 minutes at 30 °C. The mixture was then subjected to SEC using a Superose 6 column equilibrated in buffer containing 25 mM Tris-HCl pH 7.6, 150 mM NaCl, 2 mM DTT, and 5 mM MgCl_2_.

For assembly of the dCas12f-gRNA-target DNA-σ^E^-RNAP complex, a longer 99-bp target DNA, with or without mismatches in the transcription bubble, was used, consisting of the original 60-bp sequence extended by 39 bp at the 3’ end (**Supplementary Table 1**). The ternary complex was assembled following the same procedure described above, then incubated with σ^E^-RNAP overnight at 4 °C. The final complex was purified by SEC using the same equilibration buffer.

### Electron microscopy

Aliquots of 3 μL dCas12f-gRNA binary complex (1 mg/mL), dCas12f-gRNA-target DNA ternary complex (1 mg/mL), or dCas12f-gRNA-target DNA-σ^E^-RNAP complex (3 mg/mL) were applied to glow-discharged Quantifoil holey carbon girds (R1.2/1.3, 300mesh). The grids were blotted for 2.5 seconds and plunged into liquid ethane using a ThermoFisher Scientific Mark IV Vitrobot. Cryo-EM data were collected with a Titan Krios G4 microscope (FEI) operated at 300 kV and images were collected using EPU at a nominal magnification of 105,000x (resulting in a calibrated physical pixel size of 0.822 Å/pixel) with a defocus range of 0.8–2.0 μm. Three images per hole were collected using EPU with the beam-image shift method. Images were recorded on a K3 electron direct detector in super-resolution mode at the end of a GIF-Quantum energy filter operated with a slit width of 20 eV. A dose rate of 20 electrons per pixel per second and an exposure time of 2.01 seconds were used, generating 40 movie frames with a total dose of ∼59.2 electrons per Å^2^.

For the dCas12f-gRNA binary complex, we collected 4,149 micrographs. After screening based on CTF-estimated resolution and manual inspection, 3,982 micrographs were selected for further analysis. For the dCas12f-gRNA-DNA ternary complex, 2,521 micrographs were selected from a total of 2,652 collected images.

For the dCas12f-gRNA-DNA-σ^E^-RNAP complex with 99-bp native DNA, we collected 7,601 micrographs and selected 7,289 for further analysis. We then modified this complex by introducing a 15-bp mismatch region in the native DNA to facilitate RNAP open complex formation. For this modified com-plex, we collected 8,378 micrographs and selected 7,656 for further analysis.

### Image processing

Movie frames were initially imported to cryoSPARC^54^ live for on-the-fly processing during data collection. Movie frames were aligned by patch motion correction with a binning factor of 2. Contrast transfer function (CTF) parameters were estimated using Patch CTF. A few thousand particles were auto-picked without a template to generate 2D averages for subsequent template-based auto-picking. The auto-picked and extracted particles were screened by 2D classification to exclude false and bad particles that fall into 2D averages with poor features. A subset of the particles (100,000 particles) was used to generate initial models in cryoSPARC. All screened particles were subjected to heterogenous refinement using the initial models. Particles in each 3D class were subject to homogenous refinement.

Selected particles were used to generate a Topaz model for automated particle picking^55^ from all micrographs. The picked particles underwent sequential processing in cryoSPARC: 2D classification, *ab initio* reconstruction (2–4 classes), heterogeneous refinement, and Non-uniform Refinement. Results were manually inspected after each step to determine whether to rerun with modifications (e.g., adjusting class numbers) or proceed to the next step. In the final Non-uniform Refinement, we applied per-particle CTF correction. Local Refinement was performed when needed, such as for the Cas12f region in the dCas12f-gRNA-DNA-σ^E^-RNAP complex. Resolution was estimated using the FSC = 0.143 criterion. Directional resolution anisotropy was evaluated using the 3DFSC server^56^ with default parameters. Maps were post-processed using DeepEMhancer^57^ to reduce noise levels and used for model building and refinement. For the dCas12f-gRNA-DNA-σ^E^-RNAP complex, a composite map was generated by combining two maps (one from general refinement and one from local refinement around the Cas12f region) using the ‘*vop maximum’* command in Chimera^58^, which was then used for model refinement.

### Model building and refinement

AlphaFold3 models^59^ were used as initial models. The individual structures were manually fitted onto respective maps as a rigid-body in UCSF Chimera^58^ and manually adjusted in COOT^60^. The model was then fitted in COOT with all-molecule self-restrains to maintain H-bonds and stacking interactions between bases. Finally, refinement of the structure models against corresponding maps were performed using *phenix.real_space_refine* tool in Phenix^61^.

### Sequence analysis

Sequence alignments were performed using Clustal Omega^62^, and alignment diagrams were plotted using ESPript^63^. The secondary structure of gRNA (**Fig. 1j**) was visualized by Forna (force-directed RNA)^64^ and manually adjusted according to the cryo-EM structure.

### Structural analysis and visualization

Figures were generated using PyMOL, UCSF Chimera^58^ and ChimeraX^65^. To calculate the RMSD of two structures, the *align* command in PyMOL was used.

### RFP fluorescence assay

RFP assays were performed as described in Hoffmann *et al.*^6^. Briefly, *E. coli* BL21(*DE3*) cells encoding *sfGFP* in the chromosome were co-transformed with a pTarget plasmid, encoding *mRFP1* downstream of a 500 bp sequence cloned from the native *Fta* strain that includes dCas12f-gRNA target site, and a plasmid encoding *Fta* RNAP. In another round of transformations, a pEffector plasmid encoding the native *Fta* gRNA and *Fta* dCas12f-σ^E^ protein components (or mutants thereof) from the native strain (GenBank accession GCA_003584105.1) was delivered. *E. coli* cultures were grown by picking three individual colonies per sample and inoculating 5 ml of LB in 24-well plates, or 0.5 ml of LB in 96-well plates. Expression of *Fta* RNAP was induced by adding IPTG to a final concentration of 0.01 mM. Cell cultures were grown at 28 °C for 20-24 h. Then, 200 μl from each well was transferred to a 96-well optical bottom plate (Thermo Scientific) to take endpoint measurements for OD_600_ mRFP fluorescence using a Synergy Neo2 microplate reader (Biotek).

### Western Blot

*E. coli* BL21(DE3) strains containing either wild-type or mutant dCas12f and σ^E^ were grown overnight in LB. Overnight cultures were back diluted 1:200 and grown to an OD_600_ of 0.6–1.0. The cells were pelleted at 4,000 g for 10 min at 4 °C. The supernatant was removed, and the pellet was resuspended in 100 µL of lysis buffer (50 mM Tris pH 7–8, 300 mM NaCl, 0.1% detergent, 5% (v/v) glycerol) per each OD unit. Samples incubated at room temperature for 5–10 minutes before adding 1% (v/v) N-lauroylsarcosine and incubating again for 5–10 minutes. A 6X loading buffer (375 mM Tris-HCl, pH 6.8, 50% (w/v) glycerol, 9% SDS, 0.03% bromophenol blue) was added with DTT to a final concentration of 50 mM. Samples were boiled at 98 °C for 10 minutes and loaded onto a 4–20% polyacrylamide gel and ran for 25 minutes. After electrophoresis, samples were transferred onto a 0.2-μm polyvinylidene difluoride (PVDF) membrane. Anti-Flag, GAPDH antibody (Thermo Fisher Scientific), and horseradish peroxidase (HRP)–conjugated goat anti-mouse IgG1 antibody (Sigma-Aldrich) were used at concentrations of 1:10,000 and 1:30,000, respectively. SuperSignal West Dura Chemiluminescent Substrate (Thermo Fisher Scientific) was used for blot development. Blots were imaged using an Amersham ImageQuant 800 (Cytiva).

### Mass spectrometry

The bands of interest were excised using razor blade and a Trypsin in-gel digestion was carried out as previously described^66^. The peptides were subjected to a C18 desalting protocol and subsequently subjected to mass spectrometry analysis. The peptides were analyzed using Brucker’s TIMS TOF^HT^ (Bruker Daltonics Gmbh) mass spectrometer coupled with the nano Elute 2 (Bruker Daltonics gmbh) reverse phase liquid chromatography system coupled to a Captive Spray 2 ion source. The mass spectrometry settings for data-independent acquisition and the LC gradient were as previously described^67^.

The raw (.d) data was analyzed using Dia-NN software (v2.0)^68^ and Spectronaut (v19) (Biognosys). The library-free approach was employed. The spectral library was built using the *E. coli* proteome downloaded from the Uniprot database, supplemented with *Fta* dCas12f and the *Fta* RNAP α subunit. The MaxLFQ intensity was quantified by the Dia-NN software (v2.0).

### F. taeanensis σ phylogeny

The amino acid sequence for *E. coli* K-12 MG1655 (genomic accession NC_000913.3) σ^E^ (protein accession: WP_001295364.1) and σ^70^ (protein accession WP_000437376.1) was used to perform a BLASTp search in the *F. taenensis* genome (genomic accession GCA_003584105.1) to identify all *Fta*-encoded σ factors. To ensure that all σ^D^ and σ^E^-encoding genes were discovered, another BLASTp search using an *Fta*-encoded σ^D^ (protein accession RIV53857.1) and σ^E^ (protein accession RIV50554.1) was performed, respectively. The resulting 30 σ^D^ and σ^E^ sequences were used to build a Hidden Markov Model (HMM) using the hmmbuild command. The output file was used to perform an hmmsearch against the *Fta* proteome which yielded 46 hits, of which 34 were unique. Four sequences with non-σ annotations were manually excluded. MAFFT (LINSI option) was used to generate a multiple sequence alignment of the *Fta* σ sequences. A phylogenetic tree was generated using the following command: iqtree2 [alignment.fasta] -nt AUTO -m WAG -alrt 1000 -bb 1000 -abayes. The unrooted phylogenetic tree was generated in iTOL (online). σ homologs were chosen for gene synthesis to reflect the phylogenetic diversity of *Fta* σ factors.

### RpoZ conservation analysis

All proteins annotated as RpoZ were downloaded from the SwissProt database^69^ (UniProtKB, reviewed entries only), and were merged with a manually curated set of ten RpoZ sequences from genomes encoding dCas12f (**Supplementary Data 2)**. A multiple sequence alignment was constructed using MAFFT^70^ (L-INS-i algorithm). Uniprot sequences were then matched to sequences in the NCBI non-redundant (NR) protein database via a BLASTp^71^ search. Taxonomic information was then retrieved using the taxize R package^72^ (version 0.9.98) to obtain NCBI taxonomy UIDs, followed by classification data extraction. To reduce redundancy while preserving RpoZ representatives from dCas12f-encoding genomes, one sequence per species was selected for tree construction. A taxonomic tree was generated using the NCBI Common Tree tool^73^, and phylogenetic visualization was performed using ggtree^74^. Amino acids from the MAFFT-generated MSA were color-coded according to biochemical properties (hydrophobic, polar, basic, acidic, and aromatic residues), and displayed alongside the taxonomic tree. **Quantification and statistical analysis.** Detailed statistical information for the experiments is provided in the figure legends. Statistical validation for the final structural models was performed using Phenix^61^.

## Supporting information

Supplementary Information Guide

## ACKNOWLEDGMENTS

We thank Frank Vago and Thomas Klose for assistance in cryo-EM data collection, and Steven Wilson for computation. We are grateful to John Tesmer for providing equipment for radioactive experiments, and Mark Hall for helpful discussions on mass spectrometry data. The authors thank the National Institutes of Health for an S10 Award (1S10OD032364-01A1) which supported the purchase of Brucker’s TIMS-TOF^HT^, and the Purdue Proteomics Facility and its staff for assistance. This work made use of the Purdue Cryo-EM Facility. L.C. was supported by the NIH grants R01GM138675 and R35GM158248, and the NSF Faculty Early Career Development Program (CAREER) Award 2339799. S.H.S. was supported by NSF CAREER Award 2239685, a Pew Biomedical Scholarship, an Irma T. Hirschl Career Scientist Award, the Howard Hughes Medical Institute Investigator Program, and a generous startup package from the Columbia University Irving Medical Center Dean’s Office and the Vagelos Precision Medicine Fund.

## AUTHOR CONTRIBUTIONS

R.X. prepared the samples for cryo-EM and performed all biochemical assays. F.T.H and A.I.P. performed the cellular transcription assays for protein mutants. T.W. performed bioinformatics analyses. D.X. contributed to molecular cloning and protein purification. R.X. and L.C. collected cryo-EM data. L.C. and R.X. processed cryo-EM data and built the structural models. S.H.S. contributed to data analysis, interpretation, and manuscript editing. All authors participated in data analysis and contributed to the preparation of the manuscript. L.C. and S.H.S. supervised the research.

## Data availability

Cryo-EM reconstructions have been deposited in the Electron Microscopy Data Bank under the accession numbers EMD-49165, EMD-49173, EMD-49174, EMD-49175, and EMD-49176. Coordinates for atomic models have been deposited in the Protein Data Bank under the accession numbers 9N9C, 9N9M, 9N9O, 9N9P, and 9N9Q. The raw mass spectrometry data have been submitted to MassIVE repository under the identifier MSV000099155. Protein accession numbers for the *F. taeanensis* σ factor phylogeny are presented in **Extended Data** Fig. 1. Accession numbers for the RpoZ conservation analysis are listed in **Supplementary Data 2.**

## REFERENCES

1 Wiegand, T. et al. TnpB homologues exapted from transposons are RNA-guided transcription factors. Nature 631, 439–448 (2024). 10.1038/s41586-024-07598-4

2 Qi, L. S. et al. Repurposing CRISPR as an RNA-guided platform for sequence-specific control of gene expression. Cell 152, 1173–1183 (2013). 10.1016/j.cell.2013.02.022

3 Bikard, D. et al. Programmable repression and activation of bacterial gene expression using an engineered CRISPR-Cas system. Nucleic Acids Res 41, 7429–7437 (2013). 10.1093/nar/gkt520

4 Workman, R. E. et al. A natural single-guide RNA repurposes Cas9 to autoregulate CRISPR-Cas expression. Cell 184, 675–688 e619 (2021). 10.1016/j.cell.2020.12.017

5 Wu, W. Y. et al. The miniature CRISPR-Cas12m effector binds DNA to block transcription. Mol Cell 82, 4487–4502 e4487 (2022). 10.1016/j.molcel.2022.11.003

6 Hoffmann, F. T., et al. Exapted CRISPR-Cas12f homologs drive RNA-guided transcription. Submitted (2025).

7 deHaseth, P. L., Lohman, T. M., Burgess, R. R. & Record, M. T., Jr. Nonspecific interactions of Escherichia coli RNA polymerase with native and denatured DNA: differences in the binding behavior of core and holoenzyme. Biochemistry 17, 1612–1622 (1978). 10.1021/bi00602a006

8 Lee, D. J., Minchin, S. D. & Busby, S. J. Activating transcription in bacteria. Annu Rev Microbiol 66, 125–152 (2012). 10.1146/annurev-micro-092611-150012

9 Feklistov, A., Sharon, B. D., Darst, S. A. & Gross, C. A. Bacterial sigma factors: a historical, structural, and genomic perspective. Annu Rev Microbiol 68, 357–376 (2014). 10.1146/annurev-micro-092412-155737

10 Paget, M. S. & Helmann, J. D. The sigma70 family of sigma factors. Genome Biol 4, 203 (2003). 10.1186/gb-2003-4-1-203

11 Campbell, E. A. et al. Crystal structure of Escherichia coli sigmaE with the cytoplasmic domain of its anti-sigma RseA. Mol Cell 11, 1067–1078 (2003). 10.1016/s1097-2765(03)00148-5

12 Zhang, Y. et al. Structural basis of transcription initiation. Science 338, 1076–1080 (2012). 10.1126/science.1227786

13 Casas-Pastor, D. et al. Expansion and re-classification of the extracytoplasmic function (ECF) sigma factor family. Nucleic Acids Res 49, 986–1005 (2021). 10.1093/nar/gkaa1229

14 Mascher, T. Past, Present, and Future of Extracytoplasmic Function sigma Factors: Distribution and Regulatory Diversity of the Third Pillar of Bacterial Signal Transduction. Annu Rev Microbiol 77, 625–644 (2023). 10.1146/annurev-micro-032221-024032

15 Altae-Tran, H. et al. Diversity, evolution, and classification of the RNA-guided nucleases TnpB and Cas12. Proc Natl Acad Sci U S A 120, e2308224120 (2023). 10.1073/pnas.2308224120

16 Sampson, T. R., Saroj, S. D., Llewellyn, A. C., Tzeng, Y. L. & Weiss, D. S. A CRISPR/Cas system mediates bacterial innate immune evasion and virulence. Nature 497, 254–257 (2013). 10.1038/nature12048

17 Huang, C. J., Adler, B. A. & Doudna, J. A. A naturally DNase-free CRISPR-Cas12c enzyme silences gene expression. Mol Cell 82, 2148–2160 e2144 (2022). 10.1016/j.mol-cel.2022.04.020

18 Gilbert, L. A. et al. Genome-Scale CRISPR-Mediated Control of Gene Repression and Activation. Cell 159, 647–661 (2014). 10.1016/j.cell.2014.09.029

19 Takeda, S. N. et al. Structure of the miniature type V-F CRISPR-Cas effector enzyme. Mol Cell 81, 558–570 e553 (2021). 10.1016/j.molcel.2020.11.035

20 Xiao, R., Li, Z., Wang, S., Han, R. & Chang, L. Structural basis for substrate recognition and cleavage by the dimerization-dependent CRISPR-Cas12f nuclease. Nucleic Acids Res 49, 4120–4128 (2021). 10.1093/nar/gkab179

21 Hino, T. et al. An AsCas12f-based compact genome-editing tool derived by deep mutational scanning and structural analysis. Cell 186, 4920–4935 e4923 (2023). 10.1016/j.cell.2023.08.031

22 Wu, T. et al. An engineered hypercompact CRISPR-Cas12f system with boosted gene-editing activity. Nat Chem Biol 19, 1384–1393 (2023). 10.1038/s41589-023-01380-9

23 Harrington, L. B. et al. Programmed DNA destruction by miniature CRISPR-Cas14 enzymes. Science 362, 839–842 (2018). 10.1126/science.aav4294

24 Yang, H., Gao, P., Rajashankar, K. R. & Patel, D. J. PAM-Dependent Target DNA Recognition and Cleavage by C2c1 CRISPR-Cas Endonuclease. Cell 167, 1814–1828 e1812 (2016). 10.1016/j.cell.2016.11.053

25 Zhang, H., Li, Z., Xiao, R. & Chang, L. Mechanisms for target recognition and cleavage by the Cas12i RNA-guided endonuclease. Nat Struct Mol Biol 27, 1069–1076 (2020). 10.1038/s41594-020-0499-0

26 Sternberg, S. H., Redding, S., Jinek, M., Greene, E. C. & Doudna, J. A. DNA interrogation by the CRISPR RNA-guided endonuclease Cas9. Nature 507, 62–67 (2014). 10.1038/nature13011

27 Liu, B., Hong, C., Huang, R. K., Yu, Z. & Steitz, T. A. Structural basis of bacterial transcription activation. Science 358, 947–951 (2017). 10.1126/science.aao1923

28 Chen, J. et al. E. coli TraR allosterically regulates transcription initiation by altering RNA polymerase conformation. Elife 8 (2019). 10.7554/eLife.49375

29 Fang, C. et al. Structures and mechanism of transcription initiation by bacterial ECF factors. Nucleic Acids Res 47, 7094–7104 (2019). 10.1093/nar/gkz470

30 Lin, W. et al. Structural basis of ECF-sigma-factor-dependent transcription initiation. Nat Commun 10, 710 (2019). 10.1038/s41467-019-08443-3

31 Li, L., Fang, C., Zhuang, N., Wang, T. & Zhang, Y. Structural basis for transcription initiation by bacterial ECF sigma factors. Nat Commun 10, 1153 (2019). 10.1038/s41467-019-09096-y

32 Bae, B., Feklistov, A., Lass-Napiorkowska, A., Landick, R. & Darst, S. A. Structure of a bacterial RNA polymerase holoenzyme open promoter complex. Elife 4 (2015). 10.7554/eLife.08504

33 Lin, W. et al. Structural Basis of Mycobacterium tuberculosis Transcription and Transcription Inhibition. Mol Cell 66, 169–179 e168 (2017). 10.1016/j.molcel.2017.03.001

34 Marin-Gonzalez, A., Vilhena, J. G., Perez, R. & Moreno-Herrero, F. A molecular view of DNA flexibility. Q Rev Biophys 54, e8 (2021). 10.1017/S0033583521000068

35 Nudler, E. RNA polymerase active center: the molecular engine of transcription. Annu Rev Biochem 78, 335–361 (2009). 10.1146/annurev.biochem.76.052705.164655

36 Geszvain, K., Gruber, T. M., Mooney, R. A., Gross, C. A. & Landick, R. A hydrophobic patch on the flap-tip helix of E.coli RNA polymerase mediates sigma(70) region 4 function. J Mol Biol 343, 569–587 (2004). 10.1016/j.jmb.2004.08.063

37 Minakhin, L. et al. Bacterial RNA polymerase subunit omega and eukaryotic RNA polymerase subunit RPB6 are sequence, structural, and functional homologs and promote RNA polymerase assembly. Proc Natl Acad Sci U S A 98, 892–897 (2001). 10.1073/pnas.98.3.892

38 Ghosh, P., Ishihama, A. & Chatterji, D. Escherichia coli RNA polymerase subunit omega and its N-terminal domain bind full-length beta’ to facilitate incorporation into the alpha2beta subassembly. Eur J Biochem 268, 4621–4627 (2001). 10.1046/j.1432-1327.2001.02381.x

39 Bu, F. et al. Cryo-EM Structure of Porphyromonas gingivalis RNA Polymerase. J Mol Biol 436, 168568 (2024). 10.1016/j.jmb.2024.168568

40 Lane, W. J. & Darst, S. A. The structural basis for promoter -35 element recognition by the group IV sigma factors. PLoS Biol 4, e269 (2006). 10.1371/journal.pbio.0040269

41 Li, J. et al. Structure of the transcription open complex of distinct sigma(I) factors. Nat Commun 14, 6455 (2023). 10.1038/s41467-023-41796-4

42 Rohs, R. et al. The role of DNA shape in protein-DNA recognition. Nature 461, 1248–1253 (2009). 10.1038/nature08473

43 Feklistov, A. & Darst, S. A. Structural basis for promoter-10 element recognition by the bacterial RNA polymerase sigma subunit. Cell 147, 1257–1269 (2011). 10.1016/j.cell.2011.10.041

44 Lim, H. M., Lee, H. J., Roy, S. & Adhya, S. A “master” in base unpairing during isomerization of a promoter upon RNA polymerase binding. Proc Natl Acad Sci U S A 98, 14849–14852 (2001). 10.1073/pnas.261517398

45 Bayley, D. P., Rocha, E. R. & Smith, C. J. Analysis of cepA and other Bacteroides fragilis genes reveals a unique promoter structure. FEMS Microbiol Lett 193, 149–154 (2000). 10.1111/j.1574-6968.2000.tb09417.x

46 Chen, S., Bagdasarian, M., Kaufman, M. G. & Walker, E. D. Characterization of strong promoters from an environmental Flavobacterium hibernum strain by using a green fluorescent protein-based reporter system. Appl Environ Microbiol 73, 1089–1100 (2007). 10.1128/AEM.01577-06

47 Feng, Y., Zhang, Y. & Ebright, R. H. Structural basis of transcription activation. Science 352, 1330–1333 (2016). 10.1126/science.aaf4417

48 Murakami, K. S., Masuda, S. & Darst, S. A. Structural basis of transcription initiation: RNA polymerase holoenzyme at 4 Å resolution. Science 296, 1280–1284 (2002). DOI 10.1126/science.1069594

49 Zuo, Y. & Steitz, T. A. Crystal structures of the E. coli transcription initiation complexes with a complete bubble. Mol Cell 58, 534–540 (2015). 10.1016/j.molcel.2015.03.010

50 Murakami, K. S., Masuda, S., Campbell, E. A., Muzzin, O. & Darst, S. A. Structural basis of transcription initiation: an RNA polymerase holoenzyme-DNA complex. Science 296, 1285–1290 (2002). 10.1126/science.1069595

51 Hubbard, B. P. et al. Continuous directed evolution of DNA-binding proteins to improve TALEN specificity. Nat Methods 12, 939–942 (2015). 10.1038/nmeth.3515

52 Feklistov, A. RNA polymerase: in search of promoters. Ann N Y Acad Sci 1293, 25–32 (2013). 10.1111/nyas.12197

53 Velappan, N., Sblattero, D., Chasteen, L., Pavlik, P. & Bradbury, A. R. M. Plasmid incompatibility: more compatible than previously thought? Protein Eng Des Sel 20, 309–313 (2007). 10.1093/protein/gzm005

54 Punjani, A., Rubinstein, J. L., Fleet, D. J. & Brubaker, M. A. cryoSPARC: algorithms for rapid unsupervised cryo-EM structure determination. Nat Methods 14, 290–296 (2017). 10.1038/nmeth.4169

55 Bepler, T. et al. Positive-unlabeled convolutional neural networks for particle picking in cryo-electron micrographs. Nat Methods 16, 1153–1160 (2019). 10.1038/s41592-019-0575-8

56 Tan, Y. Z. et al. Addressing preferred specimen orientation in single-particle cryo-EM through tilting. Nat Methods 14, 793–796 (2017). 10.1038/nmeth.4347

57 Sanchez-Garcia, R. et al. DeepEMhancer: a deep learning solution for cryo-EM volume post-processing. Commun Biol 4, 874 (2021). 10.1038/s42003-021-02399-1

58 Pettersen, E. F. et al. UCSF Chimera--a visualization system for exploratory research and analysis. J Comput Chem 25, 1605–1612 (2004). 10.1002/jcc.20084

59 Abramson, J. et al. Accurate structure prediction of biomolecular interactions with AlphaFold 3. Nature 630, 493–500 (2024). 10.1038/s41586-024-07487-w

60 Emsley, P., Lohkamp, B., Scott, W. G. & Cowtan, K. Features and development of Coot. Acta Crystallogr D Biol Crystallogr 66, 486–501 (2010). 10.1107/S0907444910007493

61 Afonine, P. V. et al. Real-space refinement in PHENIX for cryo-EM and crystallography. Acta Crystallogr D Struct Biol 74, 531–544 (2018). 10.1107/S2059798318006551

62 McWilliam, H. et al. Analysis Tool Web Services from the EMBL-EBI. Nucleic Acids Res 41, W597–600 (2013). 10.1093/nar/gkt376

63 Robert, X. & Gouet, P. Deciphering key features in protein structures with the new ENDscript server. Nucleic Acids Res 42, W320–324 (2014). 10.1093/nar/gku316

64 Kerpedjiev, P., Hammer, S. & Hofacker, I. L. Forna (force-directed RNA): Simple and effective online RNA secondary structure diagrams. Bioinformatics 31, 3377–3379 (2015). 10.1093/bioinformatics/btv372

65 Pettersen, E. F. et al. UCSF ChimeraX: Structure visualization for researchers, educators, and developers. Protein Sci 30, 70–82 (2021). 10.1002/pro.3943

66 Grantz, J. M. et al. The platelet and plasma proteome and targeted lipidome in postpartum dairy cows with elevated systemic inflammation. Sci Rep 14, 31240 (2024). 10.1038/ s41598-024-82553-x

67 Lee, Y. et al. Proteomics-based models of gene expression and cellular control of cotton fiber development. bioRxiv, 2025.2002.2005.636703 (2025). 10.1101/2025.02.05.636703

68 Demichev, V., Messner, C. B., Vernardis, S. I., Lilley, K. S. & Ralser, M. DIA-NN: neural networks and interference correction enable deep proteome coverage in high throughput. Nat Methods 17, 41–44 (2020). 10.1038/s41592-019-0638-x

69 UniProt, C. UniProt: the Universal Protein Knowledgebase in 2025. Nucleic Acids Res 53, D609–D617 (2025). 10.1093/nar/gkae1010

70 Katoh, K., Kuma, K., Toh, H. & Miyata, T. MAFFT version 5: improvement in accuracy of multiple sequence alignment. Nucleic Acids Res 33, 511–518 (2005). 10.1093/nar/gki198

71 Camacho, C. et al. BLAST+: architecture and applications. BMC Bioinformatics 10, 421 (2009). 10.1186/1471-2105-10-421

72 Chamberlain, S. A. & Szocs, E. taxize: taxonomic search and retrieval in R. F1000Res 2, 191 (2013). 10.12688/f1000research.2-191.v2

73 Schoch, C. L. et al. NCBI Taxonomy: a comprehensive update on curation, resources and tools. Database (Oxford) 2020 (2020). 10.1093/database/baaa062

74 Xu, S. et al. Ggtree: A serialized data object for visualization of a phylogenetic tree and annotation data. Imeta 1, e56 (2022). 10.1002/imt2.56

