## Extended Data Figures 1-10 for "Structural basis of RNA-guided transcription by a dCas12f-σ^E^-RNAP complex"

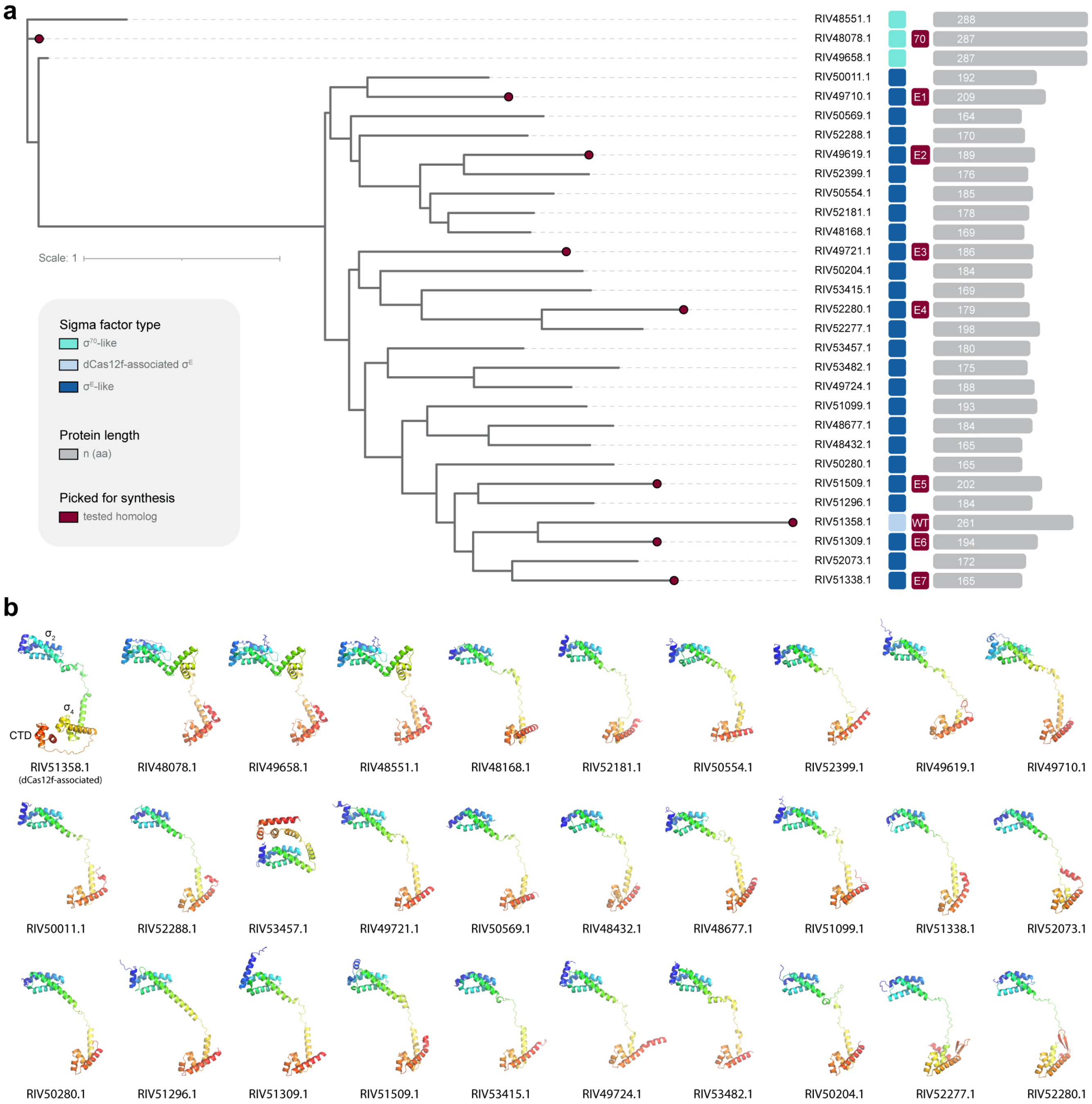


**Extended Data Figure 1 | Bioinformatics and structural analysis of all 30 σ factors in the *F. taeanensis* genome. a,** Phylogenetic analysis of the 30 σ factors, with the nine representative homologs selected for functional assays highlighted in red. **b,** AlphaFold 3 predictions of the 30 σ factors from **a**, colored in blue-to-red from the N-to-C terminus. Protein accession IDs are shown; the dCas12f-associated $\sigma^E$ (top left) possesses an additional C-terminal domain (CTD) extension.

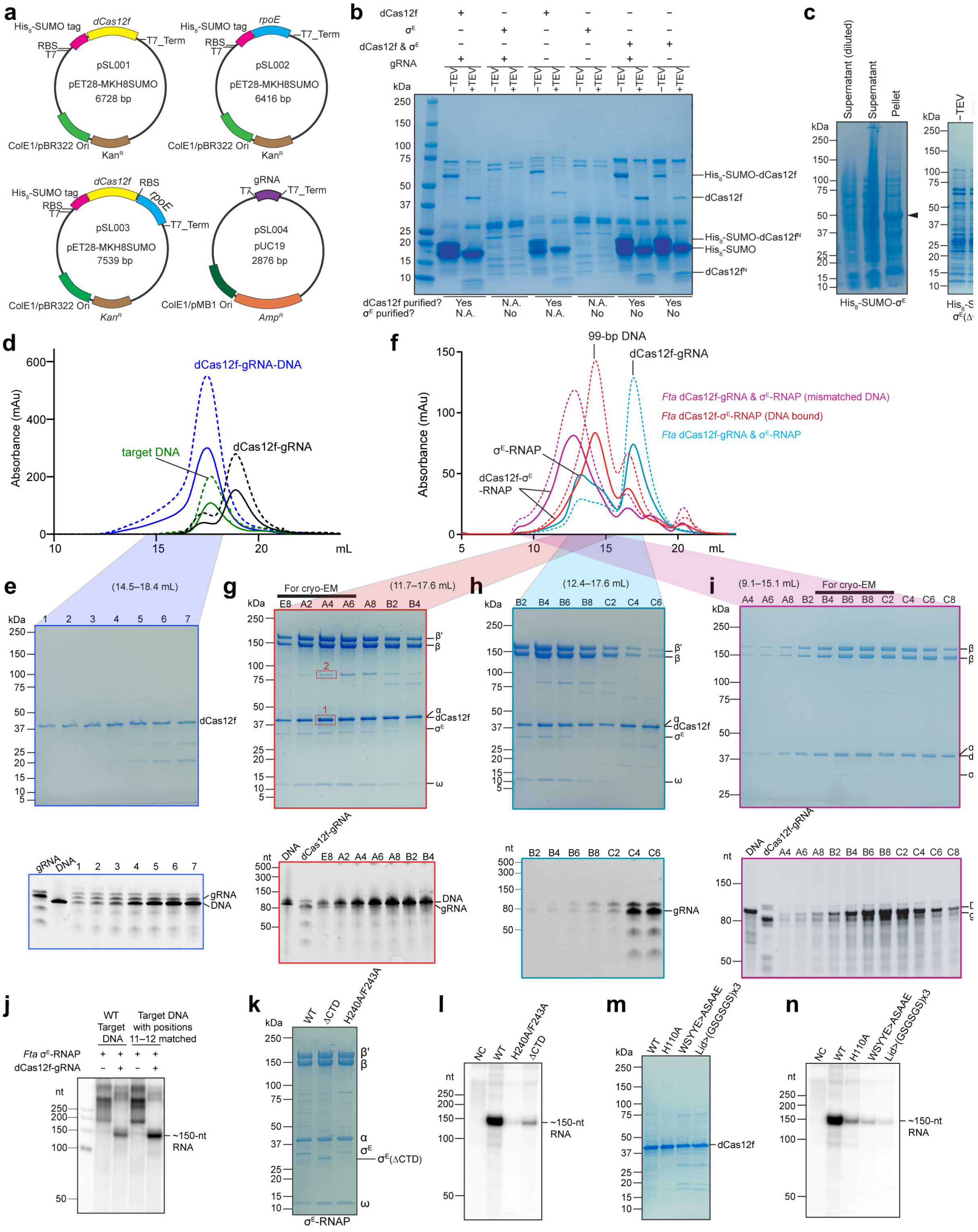


**Extended Data Figure 2 | Biochemical analysis of the dCas12f-σ$^{E}$ system. a,** Schematic of plasmid constructs used for protein expression. **b,** Purification of dCas12f and σ$^{E}$ using plasmid constructs shown in **a**. Various co-expression combinations were tested. **c,** When expressed alone, σ$^{E}$ or σ$^{E}$ with deleted CTD domain was found in the pellet rather than the supernatant, indicating expression as an inclusion body. **d,** Size exclusion chromatography (SEC) profiles of dCas12f-gRNA, target DNA alone, and the dCas12f-gRNA-DNA complex (top). UV absorbance curves at 280 nm and 260 nm are shown in solid and dashed lines, respectively. **e,** Corresponding SDS-PAGE and Urea-PAGE analyses (bottom) from the SEC experiment in **d** confirmed formation of the Cas12f-gRNA-DNA complex. **f,** SEC profiles illustrating interactions between dCas12f-gRNA and σ$^{E}$-RNAP in the presence and absence of target DNA. UV absorbance curves at 280 nm and 260 nm are shown in solid and dashed lines, respectively. **g,** Corresponding SDS-PAGE and Urea-PAGE analyses from the SEC experiment of *Fta* dCas12f-$\sigma^E$-RNAP (DNA bound) in **f**. Red dashed boxes indicate two bands analyzed by mass spectrometry shown in **Supplementary Data 1**. **h,** Corresponding SDS-PAGE and Urea-PAGE analyses from the SEC experiment of *Fta* dCas12f-gRNA and $\sigma^E$-RNAP in **f**. **i,** Corresponding SDS-PAGE and Urea-PAGE analyses from the SEC experiment of *Fta* dCas12f-$\sigma^E$-RNAP (mismatched DNA) in **f**. **j,** *In vitro* transcription assays with radiolabeled NTPs, comparing target DNA with mutations at positions 11–12 (in the wild-type target DNA, these positions contain mismatches with the gRNA). **k,** SDS-PAGE analysis of purified wild-type $\sigma^E$-RNAP, $\sigma^E$-RNAP with deletion of the CTD of $\sigma^E$ (ΔCTD), and $\sigma^E$-RNAP with two site mutations in $\sigma^E$ (H240A/F243A). **l,** *In vitro* transcription assays with radiolabeled NTPs, using $\sigma^E$-RNAP variants shown in **k**. **m,** SDS-PAGE analysis of purified wild-type and mutant dCas12f-gRNA. **n,** *In vitro* transcription assays with radiolabeled NTPs, using dCas12f-gRNA variants shown in **m**. Similar data were obtained from three independent preparations in panels **b, c, e, g–i, k,** and **m**, and from three biological replicates in panels **j, l,** and **n.**

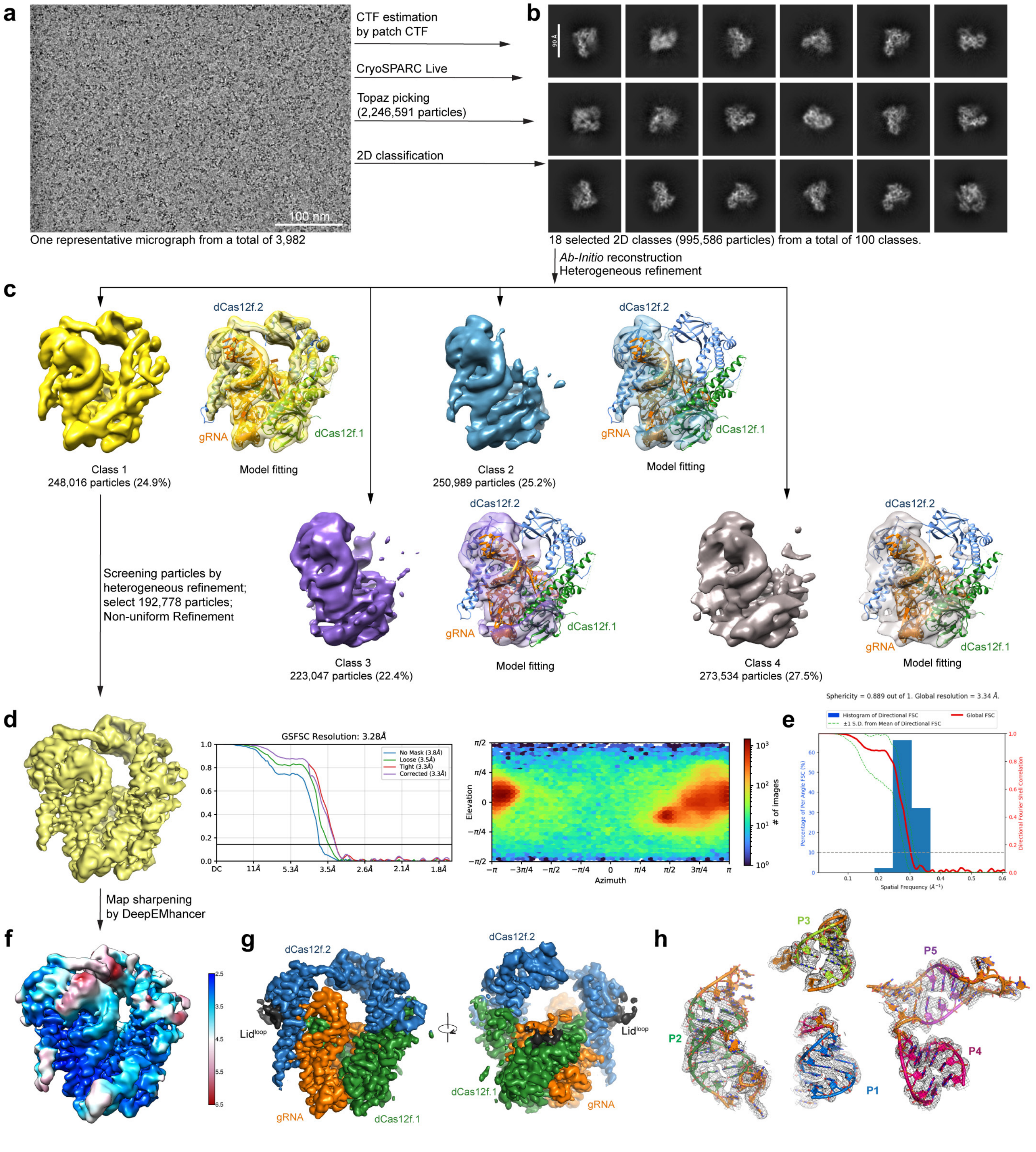


**Extended Data Figure 3 | Cryo-EM of the *Fta* dCas12f-gRNA complex. a,** A representative raw cryo-EM micrograph from a total of 3,982 micrographs. **b,** Representative 2D class averages. **c,** Four classes from heterogeneous refinement. Model fittings are shown on the right, highlighting the flexibility of the dimerization region (REC domain of dCas12f.1, and the REC and WED domains of dCas12f.2) in classes 2–4. **d,** Final non-uniform refinement of dCas12f-gRNA at 3.28 Å; global gold-standard FSC curves and particle angular distributions are shown on the right. **e,** 3D Fourier shell correlation (3DFSC) analysis of the reconstruction shown in **d**. **f,** Local resolution map. **g,** Cryo-EM map of the dCas12f-gRNA complex, after post-map sharpening by DeepEMhancer. **h,** Stem-loop structures (P1–P5) of the gRNA, with corresponding cryo-EM density shown in mesh.

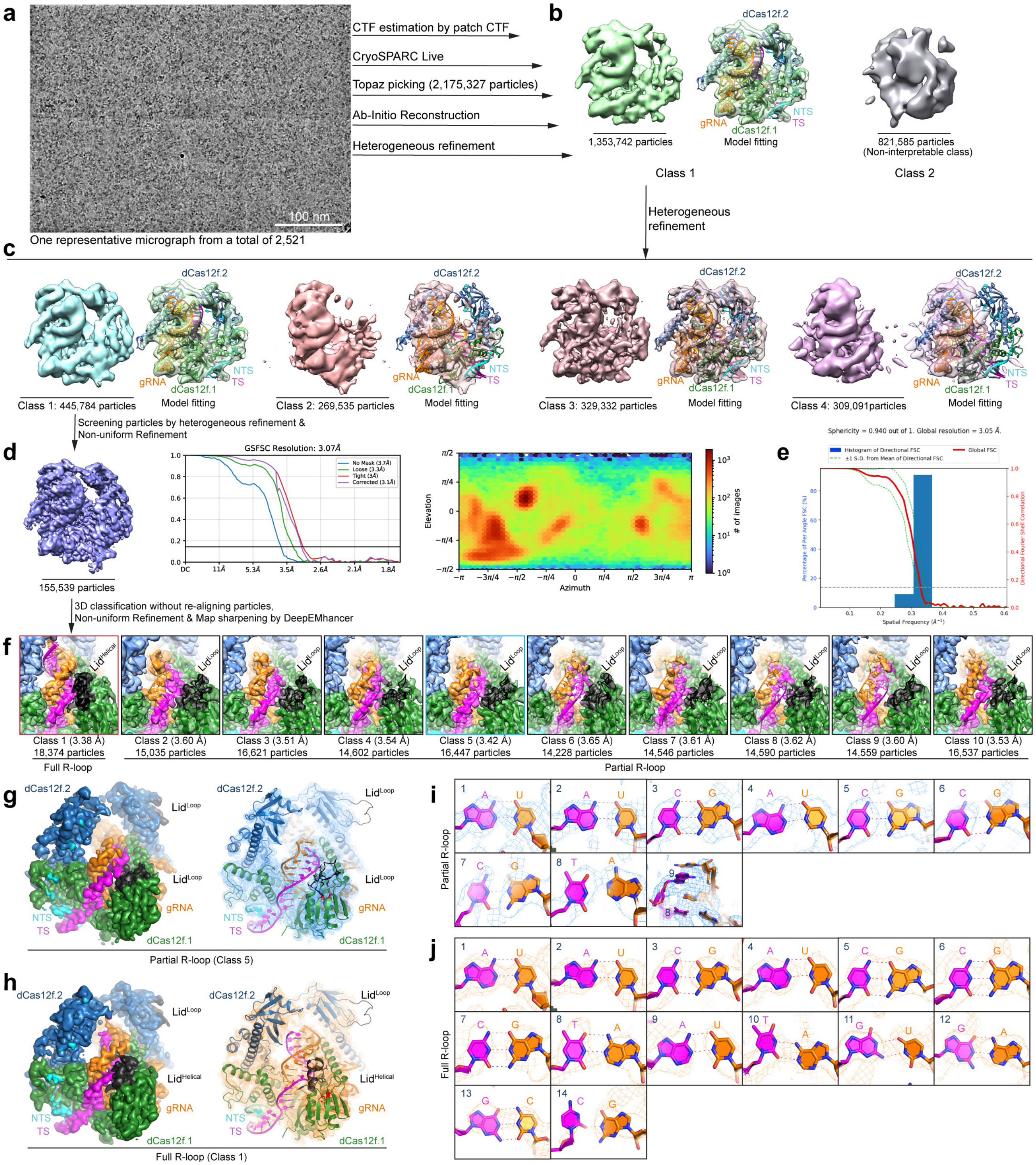


**Extended Data Figure 4 | Cryo-EM of the *Fta* dCas12f-gRNA-DNA complex. a,** A representative raw cryo-EM micrograph from a total of 2,521 micrographs. **b,** First round of heterogeneous refinement with two classes. **c,** Second round of heterogeneous refinement with four classes; model fittings are shown on the right. **d,** Non-uniform refinement of dCas12f-gRNA-DNA at 3.07 Å; global gold-standard FSC curves and particle angular distributions are shown on the right. **e,** 3DFSC analysis of the reconstruction shown in **d**. **f,** 3D classification (10 classes) without re-aligning particles, revealing more detailed variations observed in the RNA-DNA heteroduplex. Class 1 represents a full R-loop state with the lid in a helical conformation, whereas classes 2–9 represent partial R-loop conformations with the lid motif in a loop conformation. **g,h**, Final maps of the dCas12f-gRNA-DNA complex in partial and full R-loop states. **i,j,** Structure of RNA-DNA base pairing in partial and full R-loop states, with the cryo-EM maps shown in mesh.

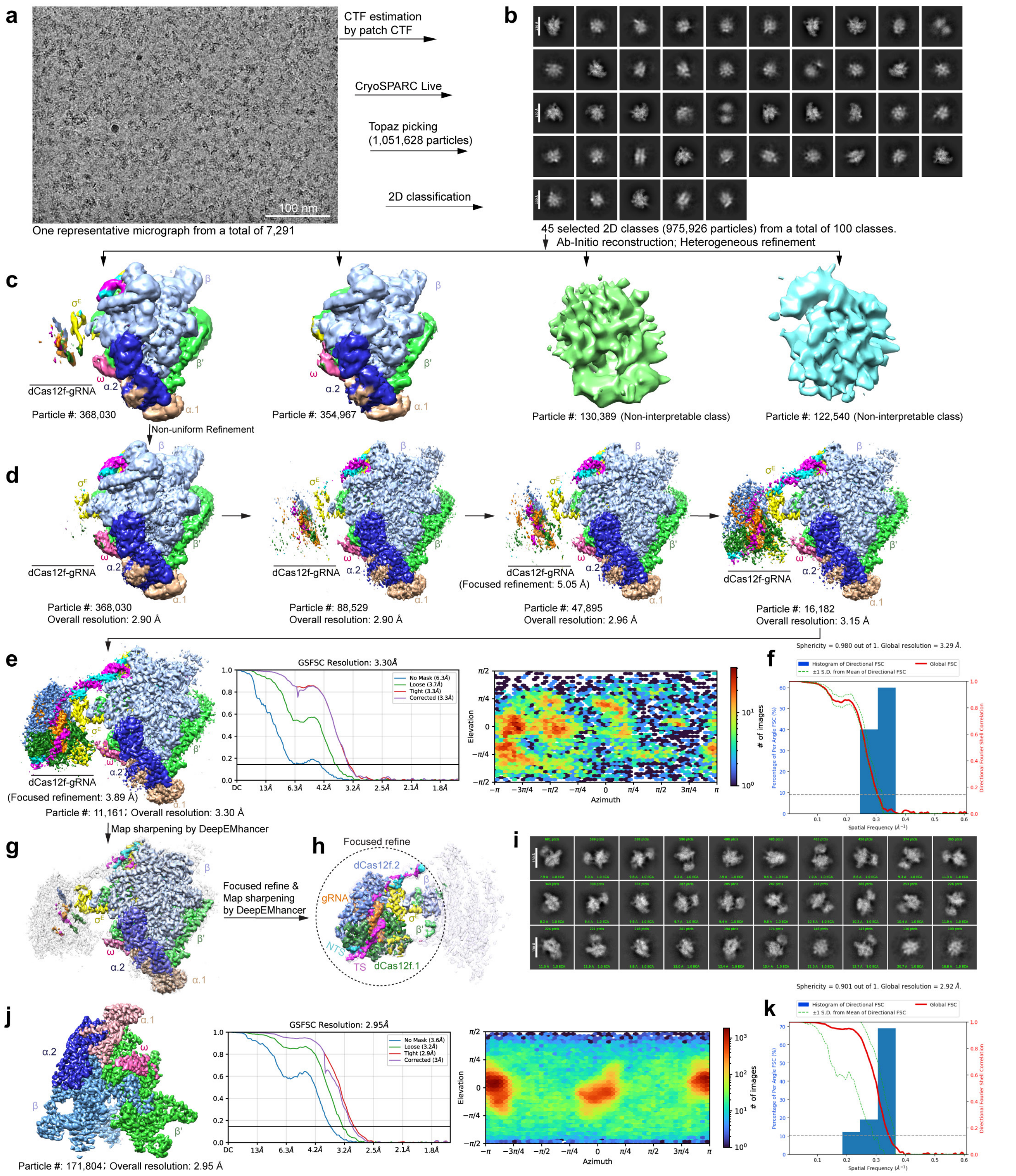


**Extended Data Figure 5 | Cryo-EM structure of the dCas12f-σ$^{E}$-RNAP complex bound to 99-bp fully matched dsDNA.** **a,** A representative raw cryo-EM micrograph from a total of 7,291 micrographs. **b,** Representative 2D class averages. **c,** Heterogeneous refinement with four classes. **d,** Particle screening through multiple rounds of heterogeneous refinement to improve the overall map of the entire complex. **e,** Non-uniform refinement from the final particle set; global gold-standard FSC curves and particle angular distributions are shown on the right. **f,** 3DFSC analysis of the reconstruction shown in **e**. **g,** Map after post-map sharpening by DeepEMhancer. **h,** Focused refinement using a mask including the dCas12f-gRNA region. **i,** Representative 2D class averages from the final particle set. **j,** Final map of apo RNAP at 2.95 Å; global gold-standard FSC curves and particle angular distributions are shown on the right. **k,** 3DFSC analysis of the reconstruction shown in **j**.

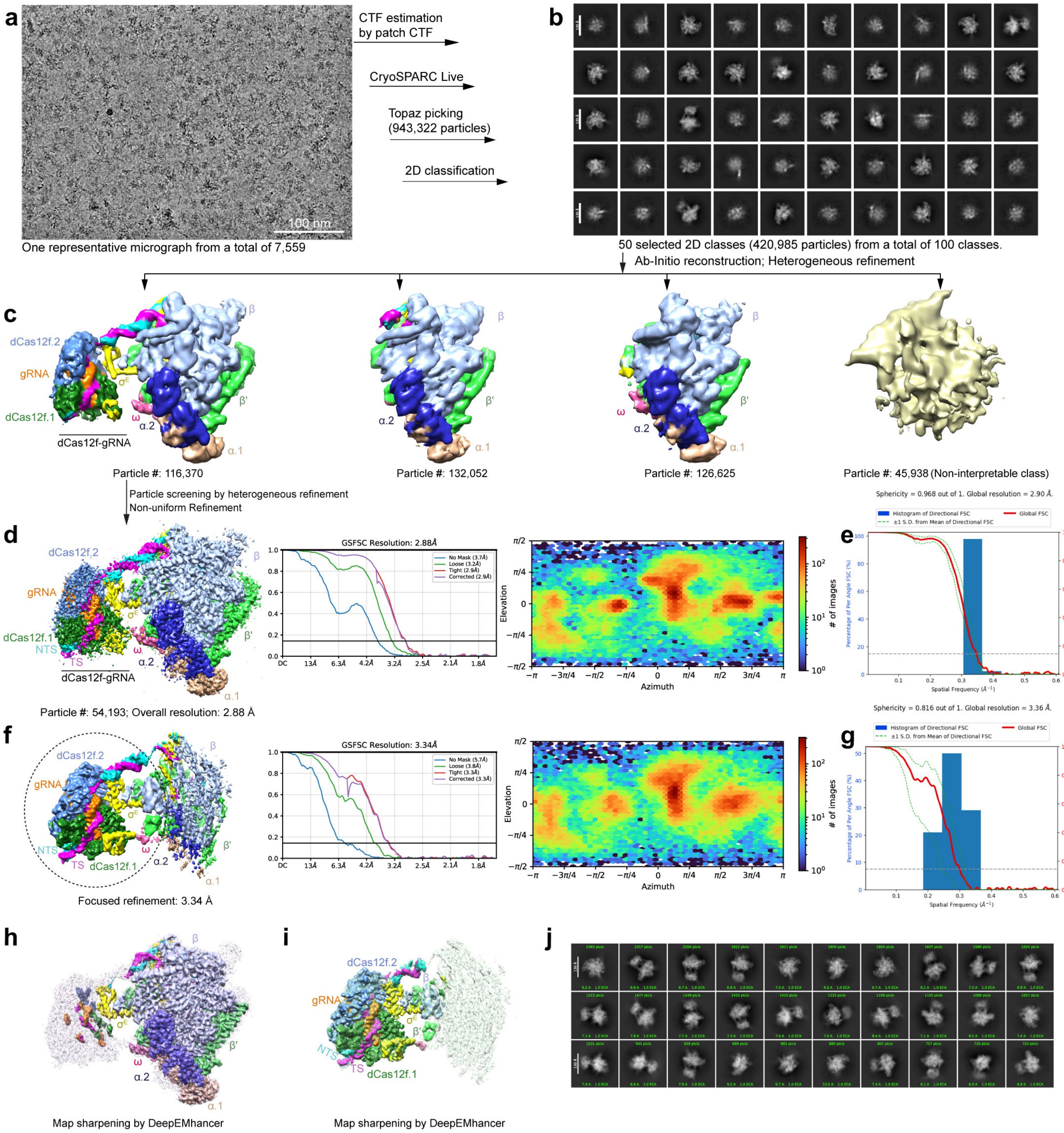


**Extended Data Figure 6 | Cryo-EM structure of the dCas12f-σ$^{E}$-RNAP complex bound to a 99-bp dsDNA containing 15-bp mismatches. a,** A representative raw cryo-EM micrograph from a total of 7,559 micrographs. **b,** Representative 2D class averages. **c,** Heterogeneous refinement with four classes. **d,** Non-uniform refinement from the final particle set; global gold-standard FSC curves and particle angular distributions are shown on the right. **e,** 3DFSC analysis of the reconstruction shown in **d**. **f,** Focused refinement using a mask including the dCas12f-gRNA region. ; global gold-standard FSC curves and particle angular distributions are shown on the right. **g,** 3DFSC analysis of the reconstruction shown in **f**. **h,** Map in **d** after post-map sharpening by DeepEMhancer. **i,** Map in **f** after post-map sharpening by DeepEMhancer. **j,** Representative 2D class averages from the final particle set.

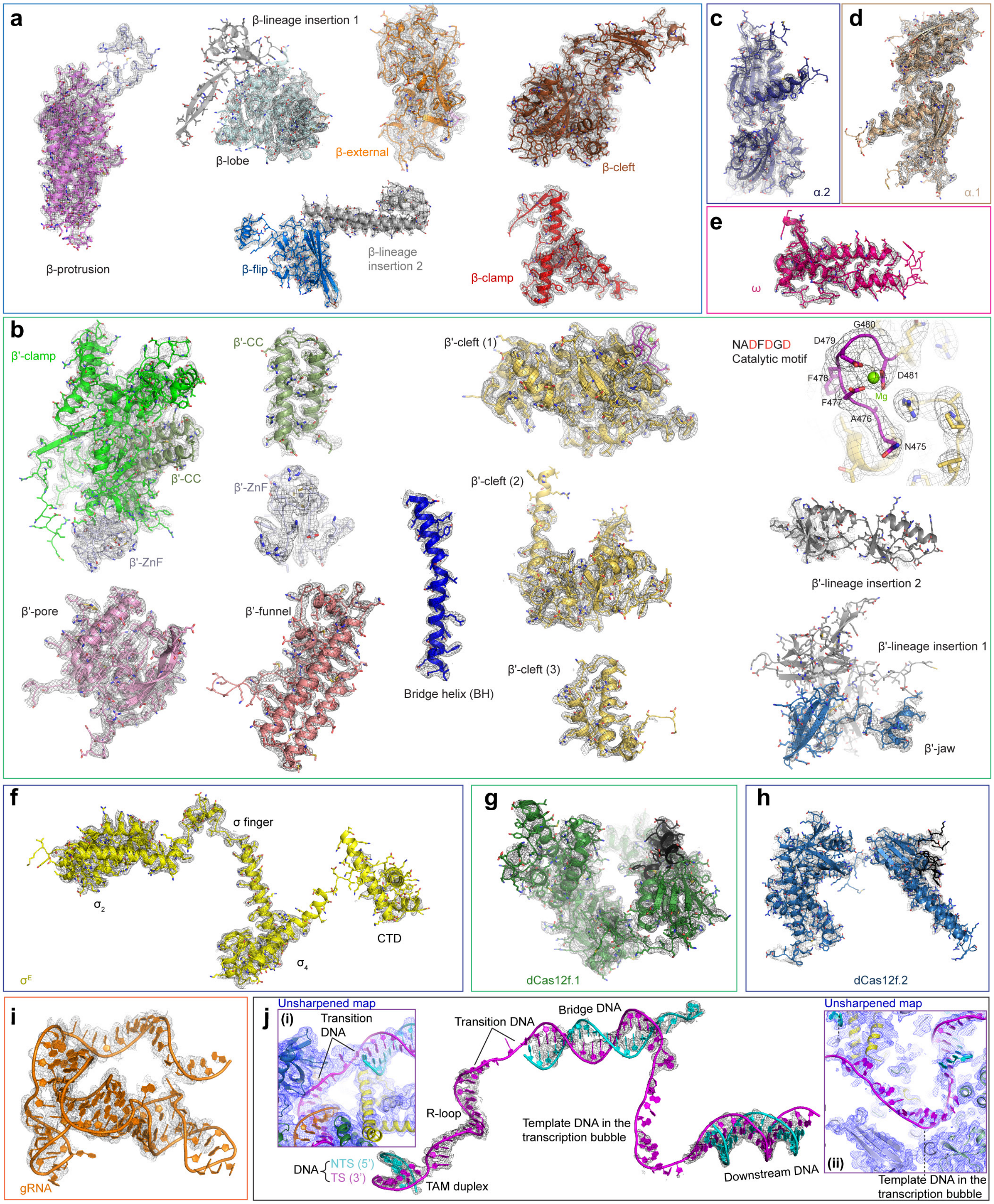


**Extended Data Figure 7 | Model-to-map fitting for each subunit of the DNA-bound dCas12f-σ$^{E}$-RNAP complex. a,** Cartoon model of each domain in β with cryo-EM density shown in mesh. **b,** Domains in β'. **c,** α.2-NTD. **d,** α.1-NTD. **e,** ω. **f,** σ$^{E}$. **g,** dCas12f.1. **h,** dCas12f.2. **i,** gRNA. **j,** Target DNA. Insets (i) and (ii) show the unsharpened map in blue mesh, where the transition DNA and transcription bubble regions are visible but not well-defined in the sharpened map.

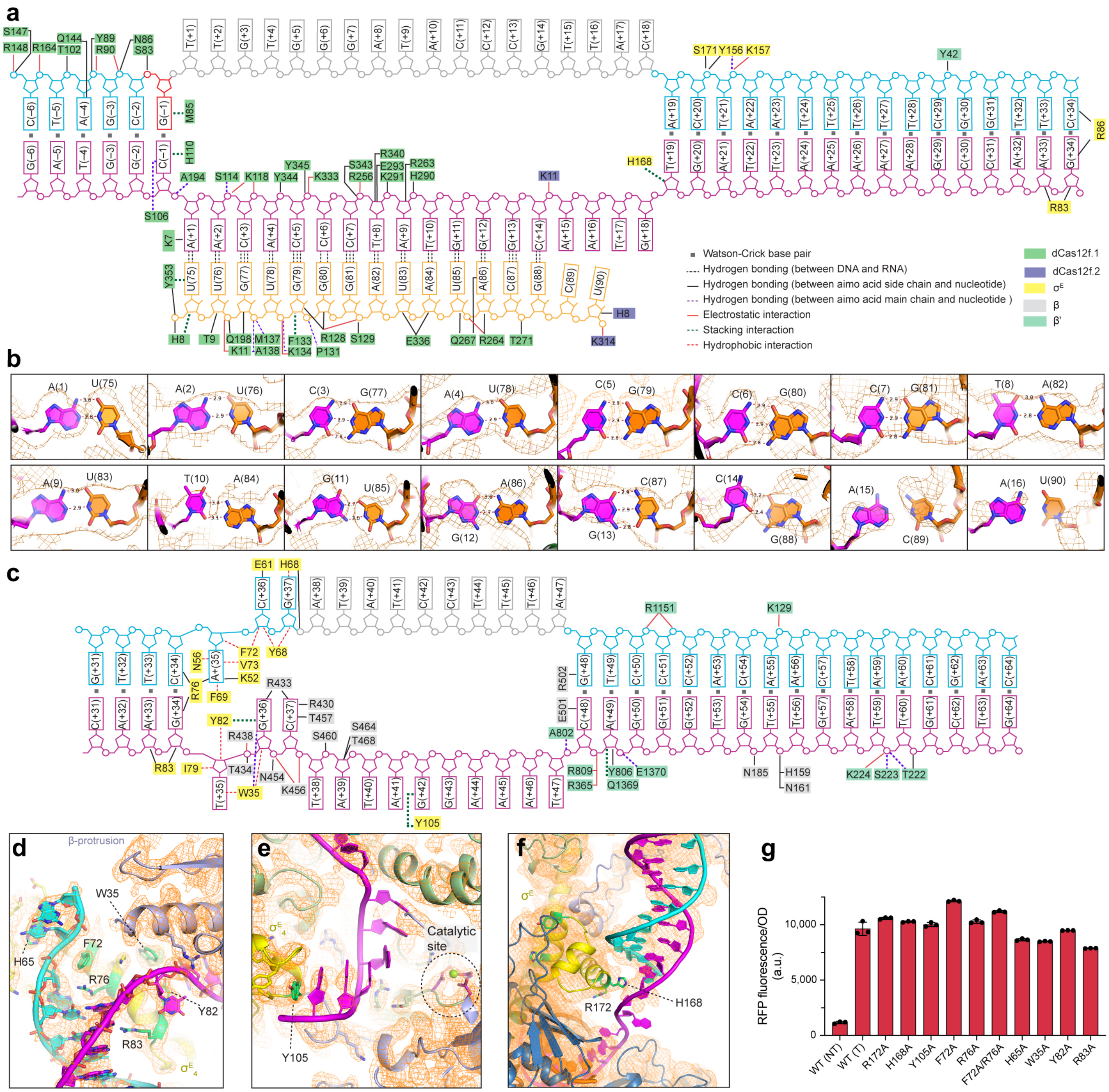

**Extended Data Figure 8 | Interaction between target DNA and protein subunits in the dCas12f-σ<sup>E</sup>-RNAP complex. a,** Diagram of the target DNA from position -6–34, and its interactions with protein subunits in the complex. **b,** Structure of target DNA and gRNA from positions 1–16 and positions 75–90, respectively, with the cryo-EM map shown in mesh. **c,** Diagram of the target DNA from position 31–64, and its interactions with protein subunits in the complex. **d,** Detailed interactions between the $\sigma^E_2$ domain and the target DNA, with residues tested by alanine substitution highlighted. **e,** Target DNA (template strand) positioned at the catalytic site of RNAP. **f,** Detailed interactions between the $\sigma^E_4$ domain and the target DNA, highlighting contacts with the minor groove. **g,** OD-normalized RFP fluorescence from cellular transcriptional reporter assays using the $\sigma^E$ mutants indicated in **d–f**. RNA-guided transcription was tolerant to the single mutations tested. Data are shown as mean ± s.d. for $n$ = 3 biologically independent samples.

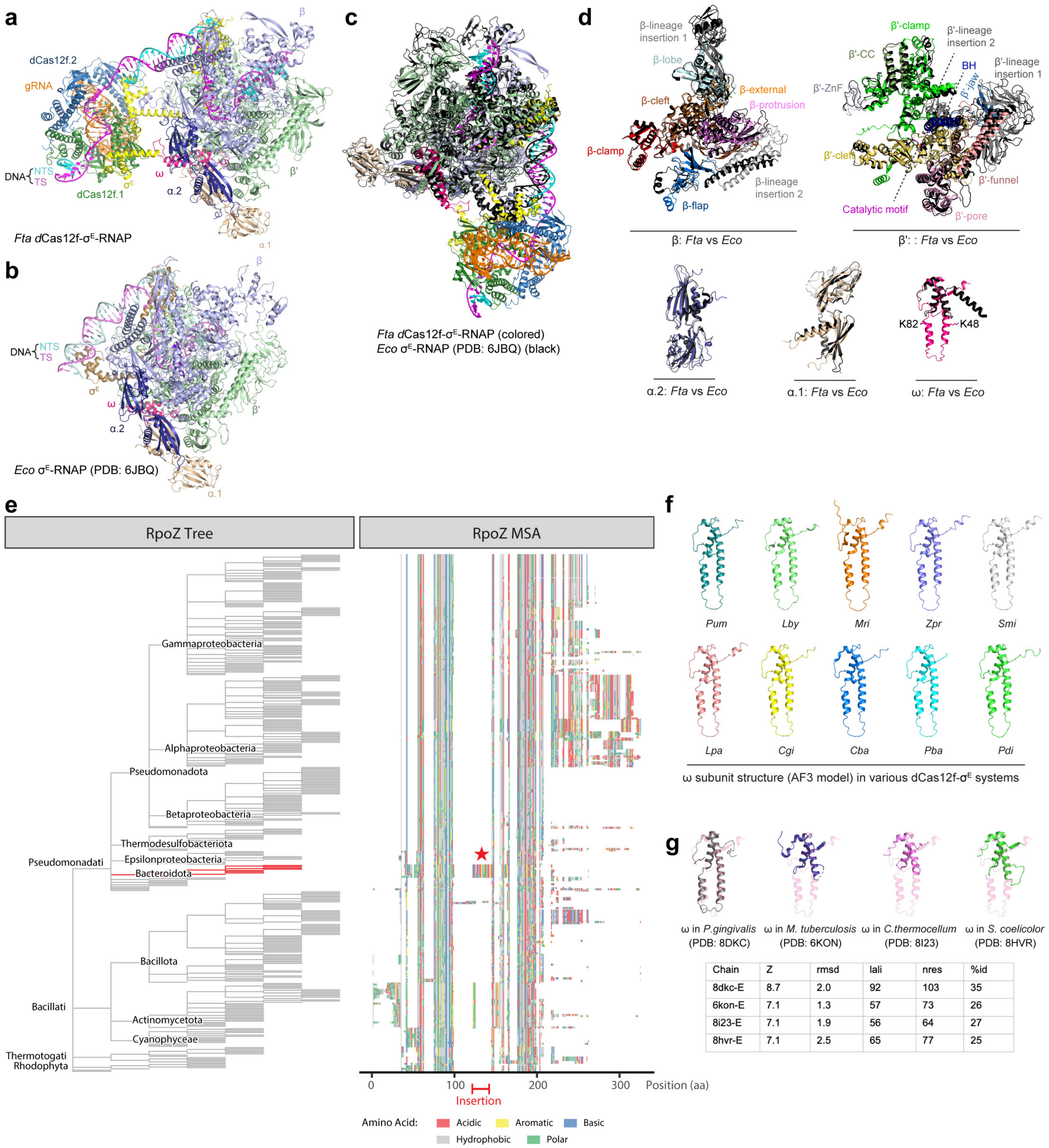


| Chain | Z | rmsd | lali | nres | %id |
|---|---|---|---|---|---|
| 8dkc-E | 8.7 | 2.0 | 92 | 103 | 35 |
| 6kon-E | 7.1 | 1.3 | 57 | 73 | 26 |
| 8i23-E | 7.1 | 1.9 | 56 | 64 | 27 |
| 8hvr-E | 7.1 | 2.5 | 65 | 77 | 25 |

**Extended Data Figure 9 | Structural comparison of DNA-bound *Fta* dCas12f-σ$^E$-RNAP and *E. coli* σ$^E$-RNAP complexes.** **a,** Structure of DNA-bound dCas12f-σ$^E$-RNAP. **b,** Structure of DNA-bound *Eco* σ$^E$-RNAP. **c,** Superimposed structures. **d,** Superimposed structures of individual RNAP core subunit. **e,** A phylogenetic tree and accompanying multiple sequence alignment of all bacterial RpoZ sequences in the SwissProt database, merged with ten unique RpoZ sequences from dCas12f-encoding genomes. **f,** AlphaFold structures of representative ω subunit from genomes containing dCas12f-σ$^E$ systems. **g,** Top DALI search results using *Fta* ω as a query; *Fta* ω is shown in transparent for comparison.

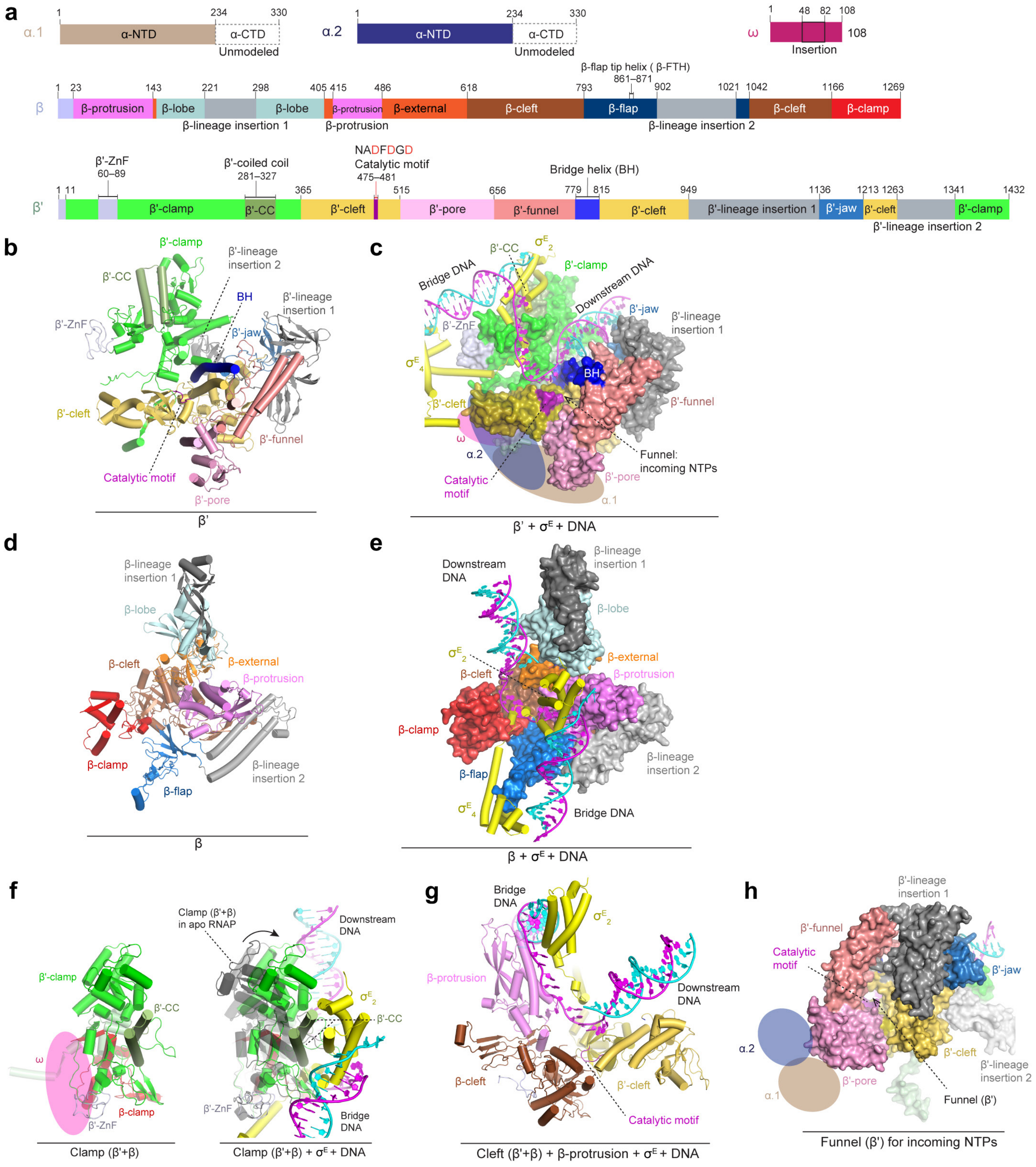

**Extended Data Figure 10 | Structure of *Fta* RNAP core subunits. a,** Domain organization of the RNAP core subunits. **b,** Structure of the β' subunit with each domain color-coded. **c,** Structure of β' in the context of $\sigma^E$ and DNA. **d,** Structure of the β subunits with each domain color-coded. **e,** Structure of β in the context of $\sigma^E$ and DNA. **f,** Structure of the Clamp domain alone (left) and in the context of $\sigma^E$ and DNA (right); the conformational change between both states is indicated. **g,** Structure of the 'Cleft' formed by the β-cleft, β'-cleft, and β-protrusion. **h,** View of the 'Funnel' structure formed by the β'-pore, β'-funnel, and β'-cleft domains.
