## Extended Data Table 1 for "Structural basis of RNA-guided transcription by a dCas12f-σ^E^-RNAP complex"

### Cryo-EM data collection, refinement and validation statistics

| | dCas12f-gRNA (EMDB-49165) (PDB 9N9C) | dCas12f-gRNA -DNA, Partial R-loop (EMDB-49173) (PDB 9N9M) | dCas12f-gRNA -DNA, Full R-loop (EMDB-49174) (PDB 9N9O) | RNAP (EMDB-4917 5) (PDB 9N9P) | dCas12f-$\sigma^E$-R NAP (EMDB-49176 ) (PDB 9N9Q) |
|---|---|---|---|---|---|
| **Data collection and processing** | | | | | |
| Magnification | 105000 | 105000 | 105000 | 105000 | 105000 |
| Voltage (kV) | 300 | 300 | 300 | 300 | 300 |
| Electron exposure (e–/Å$^2$) | 59.2 | 59.2 | 59.2 | 59.2 | 59.2 |
| Defocus range (μm) | 0.8–2.0 | 0.8–2.0 | 0.8–2.0 | 0.8–2.0 | 0.8–2.0 |
| Pixel size (Å) | 0.822 | 0.822 | 0.822 | 0.822 | 0.822 |
| Symmetry imposed | C1 | C1 | C1 | C1 | C1 |
| Initial particle images (no.) | 2,246,591 | 2,175,327 | 2,175,327 | 1,051,628 | 943,322 |
| Final particle images (no.) | 192,778 | 16,447 | 18,374 | 171,804 | 54,193 |
| Map resolution (Å) | 3.28 | 3.42 | 3.38 | 2.95 | 3.65 |
| FSC threshold | 0.143 | 0.143 | 0.143 | 0.143 | 0.143 |
| Map resolution range (Å) | 2.6—~20 | 2.7—~20 | 2.7—~20 | 2.4—~20 | 2.4—~20 |
| **Refinement** | | | | | |
| Initial model used (PDB code) | AlphaFold3 model | 9N9O | AlphaFold3 model | AlphaFold3 model | AlphaFold3 model, 9N9O |
| Model resolution (Å) | 3.4 | 3.4 | 3.3 | 3.1 | 2.9 |
| FSC threshold | 0.5 | 0.5 | 0.5 | 0.5 | 0.5 |
| Model resolution range (Å) | 2.6–50 | 2.7–50 | 2.7–50 | 2.4–50 | 2.4–50 |
| Map sharpening *B* factor (Å$^2$) | –145.5 | –76.6 | –83.0 | –74.0 | –62.8 |
| Model composition | | | | | |
| Non-hydrogen atoms | 7602 | 7991 | 8185 | 23174 | 38028 |
| Protein residues | 692 | 711 | 711 | 2933 | 4249 |
| Nucleotides | 88 | 100 | 109 | 0 | 197 |
| Ligands | 1 | 0 | 0 | 2 | 2 |
| *B* factors (Å$^2$) | | | | | |
| Protein | 86.65 | 100.78 | 96.03 | 74.55 | 86.71 |
| Nucleic acids | 123.80 | 90.36 | 87.47 | n.a. | 120.96 |
| Ligand | 51.09 | n.a. | n.a. | 61.21 | 82.71 |
| R.m.s. deviations | | | | | |
| Bond lengths (Å) | 0.005 | 0.004 | 0.005 | 0.003 | 0.003 |
| Bond angles (°) | 0.624 | 0.871 | 0.900 | 0.560 | 0.549 |
| Validation | | | | | |
| MolProbity score | 2.46 | 2.42 | 2.31 | 1.54 | 1.95 |
| Clashscore | 12.37 | 13.11 | 14.48 | 8.38 | 8.17 |
| Poor rotamers (%) | 3.90 | 3.48 | 3.31 | 1.30 | 2.90 |
| Ramachandran plot | | | | | |
| Favored (%) | 94.13 | 94.33 | 96.31 | 98.25 | 97.16 |
| Allowed (%) | 5.43 | 5.25 | 3.55 | 1.75 | 2.77 |
| Disallowed (%) | 0.44 | 0.43 | 0.14 | 0 | 0.07 |
