## Supplementary Information Guide for "Structural basis of RNA-guided transcription by a dCas12f-σ^E^-RNAP complex"

**Supplementary Figures**

Supplementary Figure 1 | Source data for electrophoretic separation experiments

Supplementary Figure 2 | Sequence alignment of all 30 σ factors in the *F. taeanensis* genome.

Supplementary Figure 3 | Structural comparison of *Fta* dCas12f, *As*Cas12f, and *Un*Cas12f.

Supplementary Figure 4 | Sequence alignment of representative dCas12f proteins from genomes containing dCas12f-$\sigma^E$ systems.

Supplementary Figure 5 | Western blots of dCas12f and $\sigma^E$ mutants

Supplementary Figure 6 | Sequence alignment of representative ω subunit from genomes containing dCas12f-$\sigma^E$ systems.

Supplementary Figure 7 | Sequence alignment of representative $\sigma^E$ proteins from genomes containing dCas12f-$\sigma^E$ systems.

**Supplementary Tables**

Supplementary Table 1 | Sequence of RNA and DNA substrates used in this study.

**Supplementary Data**

Supplementary Data 1 | Mass spectrometry of protein bands from Extended Data Fig. 2g. The Excel file contains eight tabs, including peptide-level outputs (Peptide_Sample 1/2), compact identification tables (Sample1/2_Peptide identification), protein-level quantification (Protein_Quantity_Sample 1/2), and MaxLFQ summaries (Sample1/2_MaxLFQ).

Supplementary Data 2 | Protein sequences used in RpoZ conservation analysis
