## Supplementary Information for "Structural basis of RNA-guided transcription by a dCas12f-σ^E^-RNAP complex"

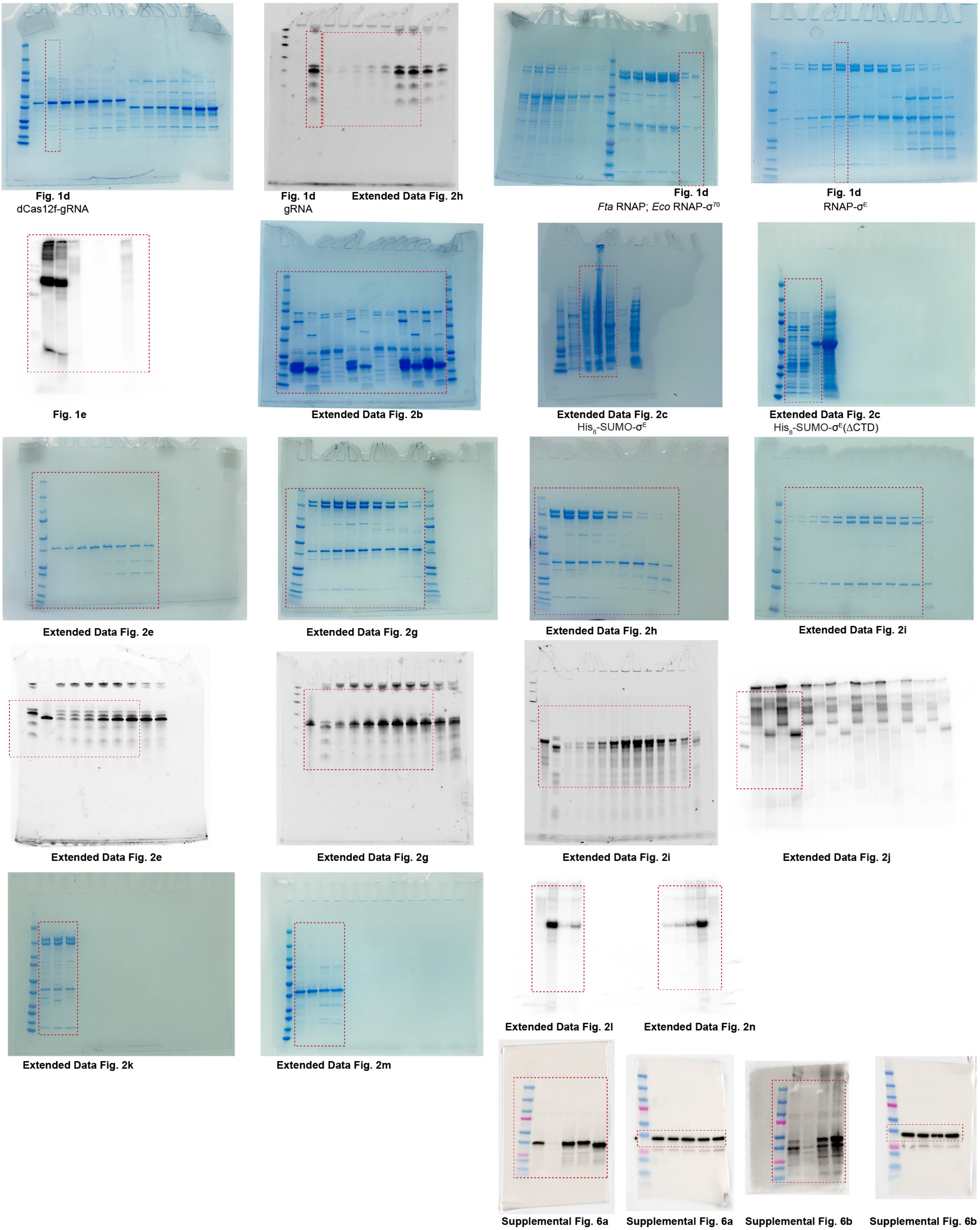


**Supplementary Figure 1 | Source data for electrophoretic separation experiments.**

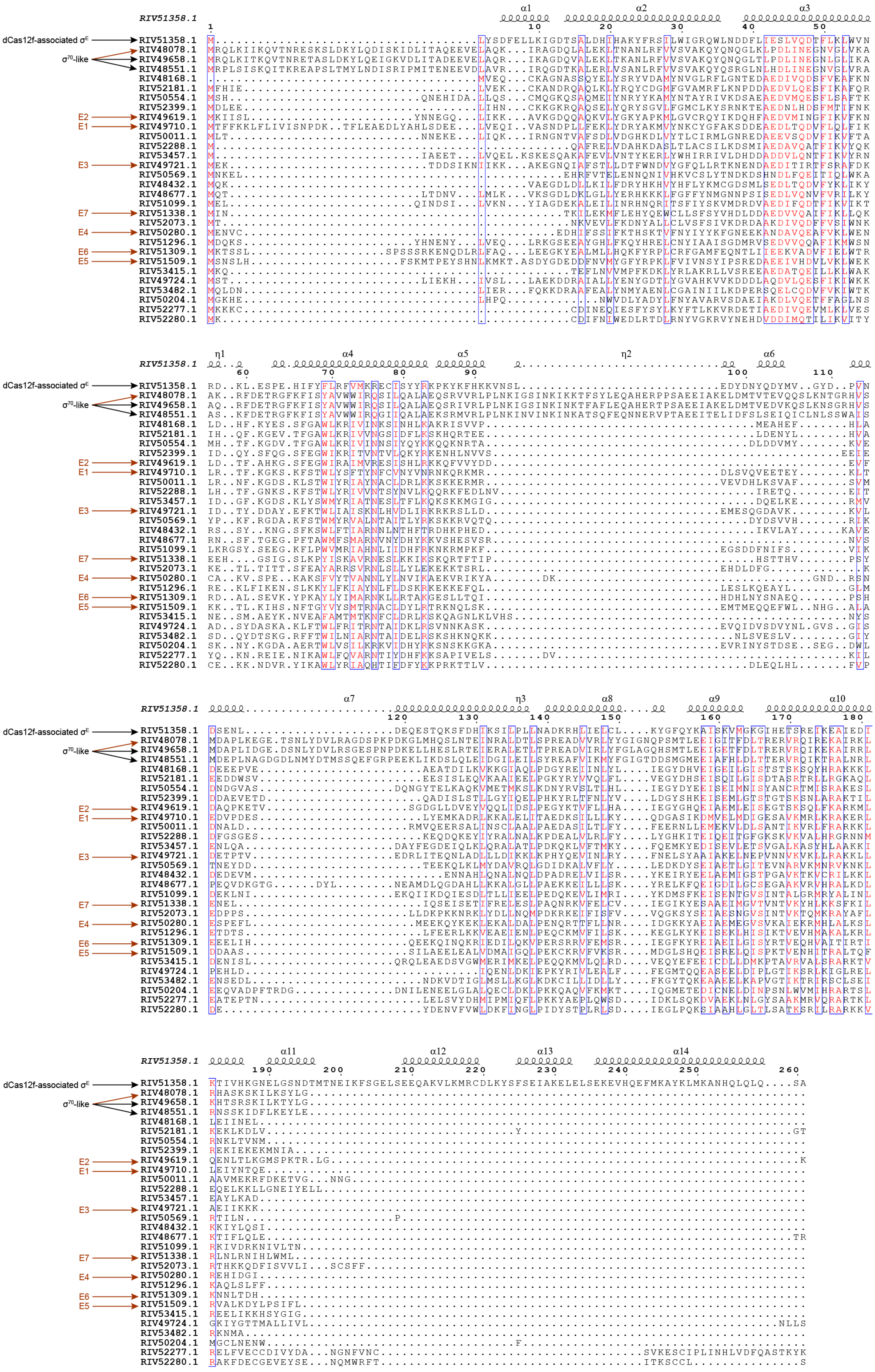


**Supplementary Figure 2 | Sequence alignment of all 30 σ factors in the F. taeanensis genome.**

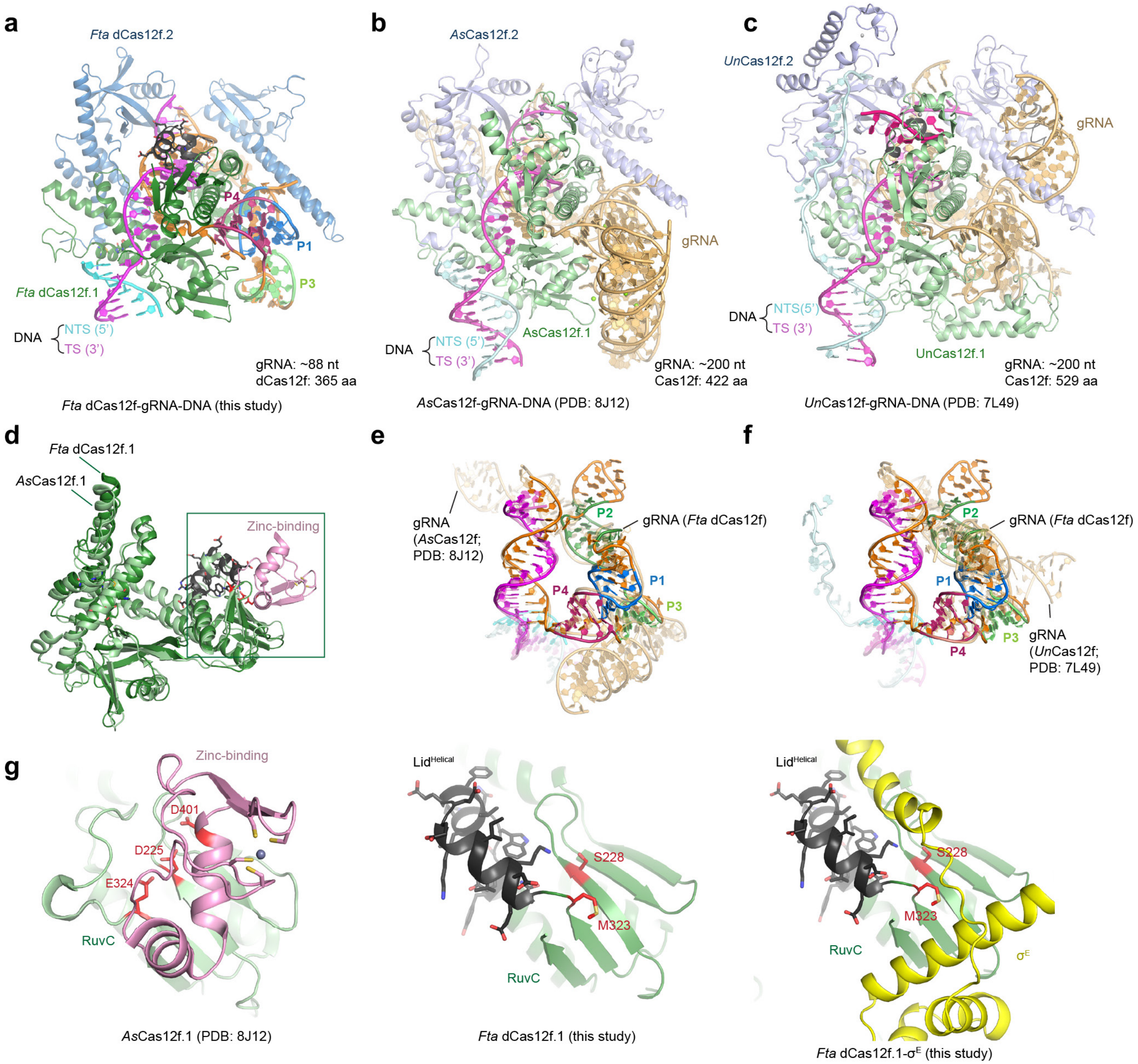


**Supplementary Figure 3 | Structural comparison of *Fta* dCas12f, *As*Cas12f, and *Un*Cas12f. a,** Structure of the *Fta* dCas12f-gRNA-DNA complex. **b,** Structure of the *As*Cas12f-gRNA-DNA complex. **c,** Structure of the *Un*Cas12f-gRNA-DNA complex. **d,** Structural alignment of *Fta* dCas12f.1 and *As*Cas12f.1. **e,** Structural alignment of gRNA structures in *Fta* dCas12f and *As*Cas12f. **f,** Structural alignment of gRNA structures in *Fta* dCas12f and *Un*Cas12f. **g,** Side-by-side comparison of the RuvC domains in *As*Cas12f.1, *Fta* dCas12f.1, and *Fta* dCas12f.1 in the DNA-bound dCas12f-$\sigma^E$-RNAP complex.

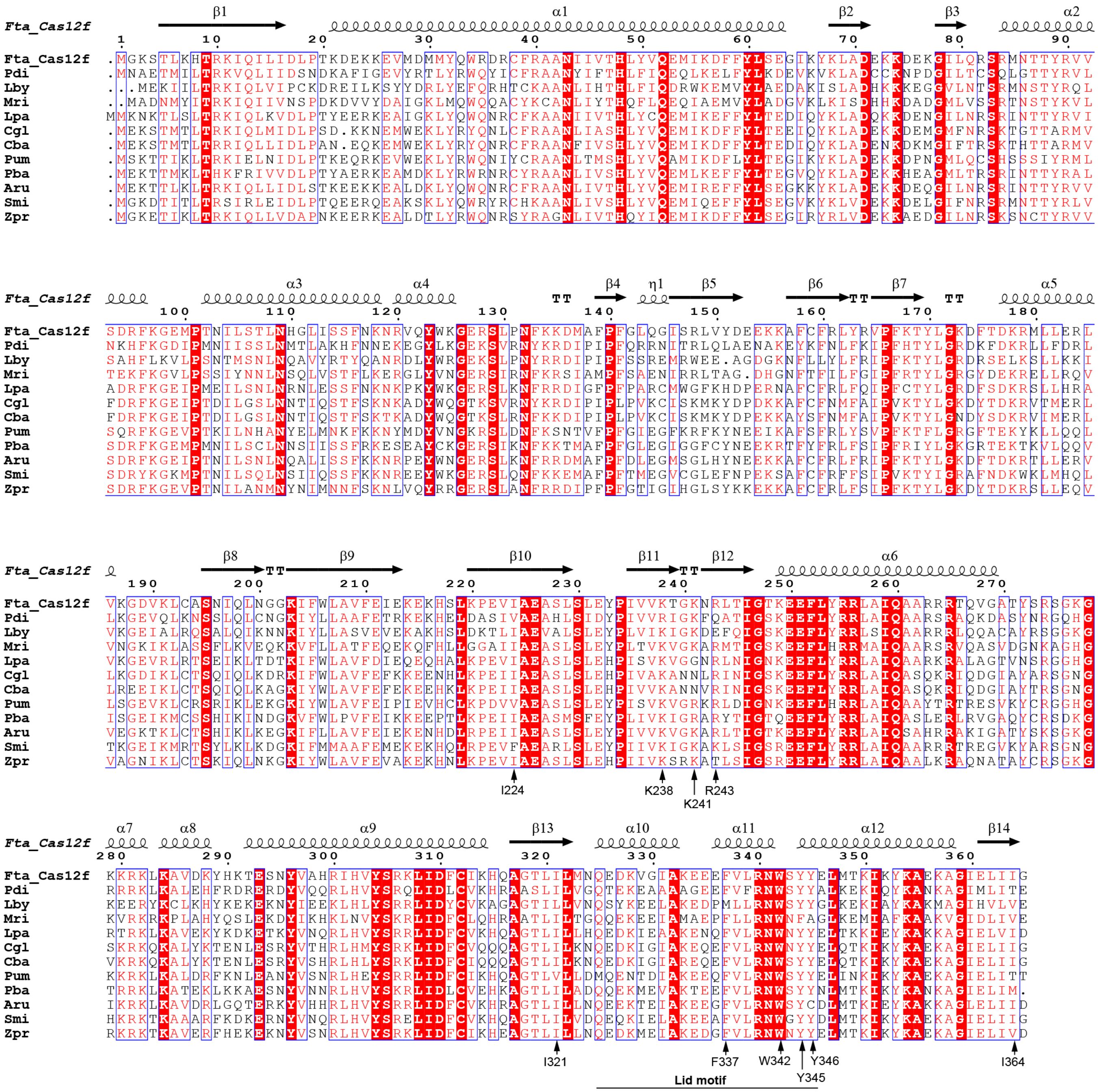


**Supplementary Figure 4 | Sequence alignment of representative dCas12f proteins from genomes containing dCas12f-σ^E systems.**

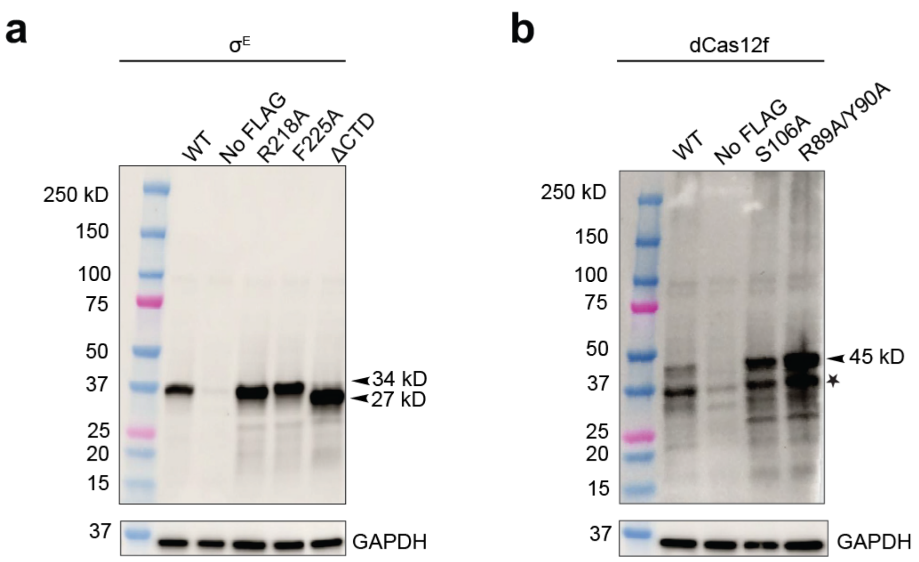


**Supplementary Figure 5 | Western blots of dCas12f and σ$^{E}$ mutants. a,** Western blot of WT σ$^{E}$ and corresponding mutants, with expected bands at 34 kD, and a band at 27 kD for the ΔCTD mutant, *n* = 2 biologically independent samples. **b,** blot of WT dCas12f and corresponding mutants with bands at expected 45 kD. An additional band marked by a star is suspected proteolysis, *n* = 2 biologically independent samples.

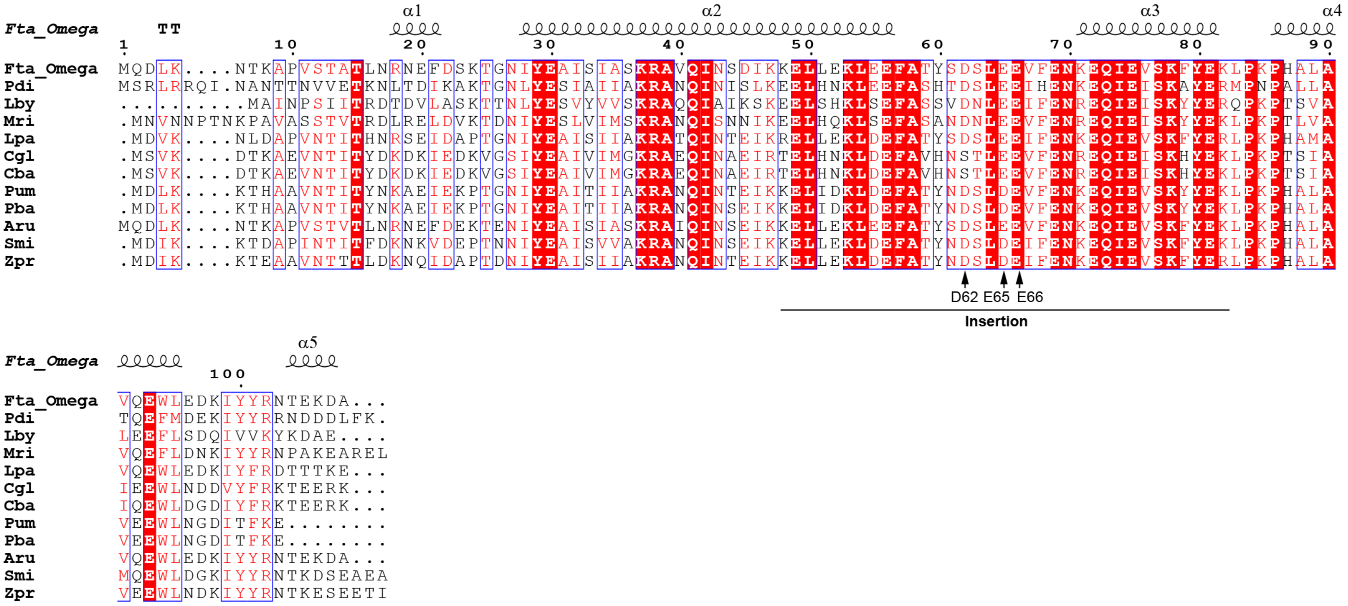


**Supplementary Figure 6 | Sequence alignment of representative ω subunit from genomes containing dCas12f-σ$^{E}$ systems.**

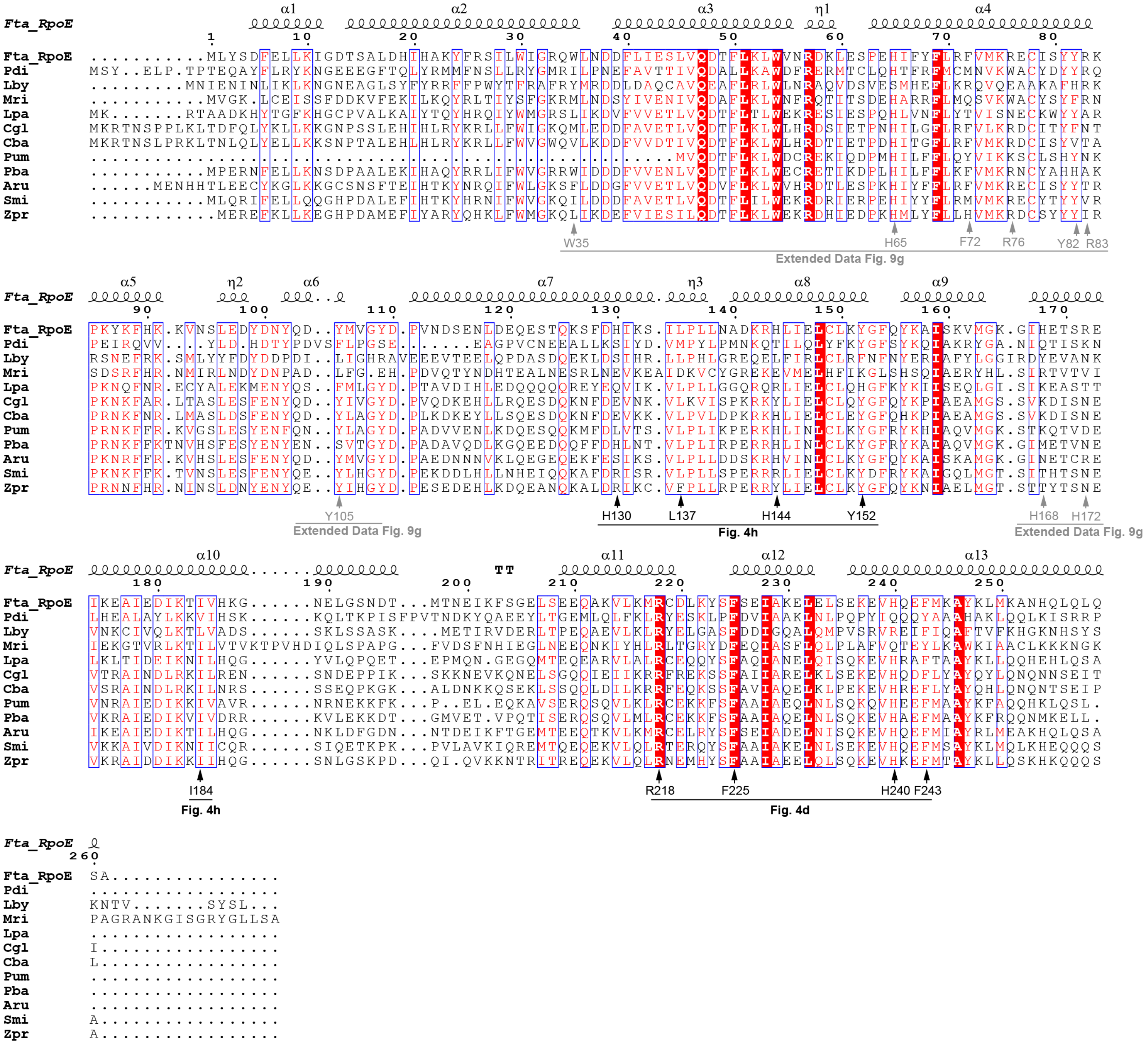


**Supplementary Figure 7 | Sequence alignment of representative σ^E proteins from genomes containing dCas12f-σ^E systems.**
