## Supplementary Table 1 for "Structural basis of RNA-guided transcription by a dCas12f-σ^E^-RNAP complex"

### Supplementary Table 1. Sequence of RNA and DNA substrates used in this study

| **RNA** | **Sequence (5'-3')** |
|---|---|
| dCas12f gRNA (native guide). Spacer region is underlined. | AUUUUGAGGGUGCUUGUACACCCAUAGGGUGAGGUUAAGAAUUACACUC<br>ACUAAGUGUGAACAACACAUACAACUUGUGGGAUAUACGCUAACA |
| dCas12f gRNA (guide for the optimized target containing an AT-rich region between TAM and TSS). Spacer region is underlined. | AUUUUGAGGGUGCUUGUACACCCAUAGGGUGAGGUUAAGAAUUACACUC<br>ACUAAGUGUGAACAACACAUACAACAUAUGCUGCAAAUCCCUGAA |
| **DNA templates for *in vitro* transcription assays** | **Sequence (5'-3')** |
| WT target DNA (Extended Data Fig. 2j). TAM is highlighted in red and TSS in blue. | TACCGAGCTCGAATTACACCCAGAAGCCGACCTCTGTTAGAGTCCAATGAA<br>GACGAATTATGTGATCTGCTTTTGCGTGCAATCGGAGAATTAAAGGACCAGG<br>TAGAGAGCGCCATCTCCACTTTTGGACTAGCGTTGTGGGATACCCGTTACAC<br>TAATTTTTCGGTTCACGATATCCTTTAGATGTATGAGCATGACTACATCGGCG<br>CCATCTTAGAGTGGGGCAAGACCAAGATCCTGTCTCCGCATATCATTAAGTAT<br>CGCGAGGACATCGAGGAGTACTGCAAAATTTCTACTAAGTGTAGATGGAAAT<br>TAGGTGCGCTTGGCAACC |
| dCas12f target DNA with positions 11–12 matched (Extended Data Fig. 2j). TAM is highlighted in red and TSS in blue. | TACCGAGCTCGAATTACACCCAGAAGCCGACCTCTGTTAGAGTCCAATGAA<br>GACGAATTATGTGATCTGCTTTTGCGTGCAATCGGAGAATTAAAGGACCAGG<br>TAGAGAGCGCCATCTCCACTTTTGGACTAGCGTTGTGGGATACtaGTTACACT<br>AATTTTTCGGTTCACGATATCCTTTAGATGTATGAGCATGACTACATCGGCGC<br>CATCTTAGAGTGGGGCAAGACCAAGATCCTGTCTCCGCATATCATTAAGTATC<br>GCGAGGACATCGAGGAGTACTGCAAAATTTCTACTAAGTGTAGATGGAAATT<br>AGGTGCGCTTGGCAACC |
| Optimized target DNA (containing an AT-rich region between TAM and TSS; used in Fig. 1e and Extended Data Fig. 2i,n). TAM is highlighted in red and TSS in blue. | TACCGAGCTCGAATTACACCCAGAAGCCGACCTCTGTTAGAGTCCAATGAA<br>GACGAATTATGTGATCTGCTTTTGCGTGCAATCGGAGAATTAAAGGACCAGG<br>TAGAGAGCGCCATCTCCACTTTTGGACTAGCGATATGCTGCAAATCCCTGAA<br>CAGCAAAAAAATGAAAAATATCAAGTAGATGTATGAGCATGACTACATCGGC<br>GCCATCTTAGAGTGGGGCAAGACCAAGATCCTGTCTCCGCATATCATTAAGT<br>ATCGCGAGGACATCGAGGAGTACTGCAAAATTTCTACTAAGTGTAGATGGAA<br>ATTAGGTGCGCTTGGCAACC |
| Non-specific DNA (used in Fig. 1e) | TACCGAGCTCGAATTACACCCAGAAGCCGACCTCTGTTAGAGTCCAATGAA<br>GACGAATTATGTGATCTGCTTTTGCGTGCAATCGGAGAATTAAAGGACCAGG<br>TAGAGAGCGCCATCTCCACTTTTGGACTAGCTTGATAAGGTAATCCAGCGAT<br>TCCCCTACAAAAACATAGCCTTTTGTAGATGTATGAGCATGACTACATCGGCG<br>CCATCTTAGAGTGGGGCAAGACCAAGATCCTGTCTCCGCATATCATTAAGTAT |

| | |
|---|---|
| | CGCGAGGACATCGAGGAGTACTGCAAAATTTCTACTAAGTGTAGATGGAAAT<br>TAGGTGCGCTTGGCAACC |
| *E coli* $\sigma^{70}$ specific target DNA (used in Fig. 1e). The –35 and –10 promoter elements are highlighted in blue and red, respectively. | TACCGAGCTCGAATTACACCCAGAAGCCGACCTCTGTTAGAGTCCAATGAA<br>GACGAATTATGTGATCTGCTTTTGCGTGCAATCGGAGAATTAAAGGACCAGG<br>TAGAGAGCGCCATCTCCACTTTTGGAcgACTTGACATCCCACCTCACGTATGC<br>TATAATGGGTGCAGATGTATGAGCATGACTACATCGGCGCCATCTTAGAGTGG<br>GGCAAGACCAAGATCCTGTCTCCGCATATCATTAAGTATCGCGAGGACATCG<br>AGGAGTACTGCAAAATTTCTACTAAGTGTAGATGGAAATTAGGTGCGCTTGG<br>CAACC |
| DNA template for 250-nt RNA marker. The T7 promoter is highlighted in green.<br>Transcribed region region is underlined. | TACCGAGCTCGAATTACACCCAGAAGCCGcgatTAATACGACTCACTATAGGAT<br>GTATGAGCATGACTACATCGGCGCCATCTTAGAGTGGGGCAAGACCAAGATC<br>CTGTCTCCGCATATCATTAAGTATCGCGAGGACATCGAGGAGTACTGCAAAC<br>CTCTGTTAGAGTCCAATGAAGACGAATTATGTGATCTGCTTTTGCGTGCAATC<br>GGAGAATTAAAGGACCAGGTAGAGAGCGCCATCTCCACTTTTGGAGATATTA<br>AATTGGTAACCAGCACTGAGTATGAGGAATTGTGGCA |
| DNA template for 200-nt RNA marker. The T7 promoter is highlighted in green.<br>Transcribed region region is underlined. | TACCGAGCTCGAATTACACGAGAGCGCCATCTCCACTTTTGGAGATATCAAA<br>TTGGTAACCAGCACTGACCAGAAGCCGcgatTAATACGACTCACTATAGGATG<br>TATGAGCATGACTACATCGGCGCCATCTTAGAGTGGGGCAAGACCAAGATCC<br>TGTCTCCGCATATCATTAAGTATCGCGAGGACATCGAGGAGTACTGCAAACC<br>TCTGTTAGAGTCCAATGAAGACGAATTATGTGATCTGCTTTTGCGTGCAATCG<br>GAGAATTAAAGGACCAGGTAGTATGAGGAATTGTGGCA |
| DNA template for 150-nt RNA marker. The T7 promoter is highlighted in green.<br>Transcribed region region is underlined. | TACCGAGCTCGAATTACACGAGAGCGCCATCTCCACTTTTGGAGATATcAAAT<br>TGGTAACCAGCACTGAAATTATGTGATCTGCTTTTGCGTGCAATCGGAGAAT<br>TAAAGGACCAGGTACCAGAAGCCGcgatTAATACGACTCACTATAGGATGTATG<br>AGCATGACTACATCGGCGCCATCTTAGAGTGGGGCAAGACCAAGATCCTGTC<br>TCCGCATATCATTAAGTATCGCGAGGACATCGAGGAGTACTGCAAACCTCTG<br>TTAGAGTCCAATGAAGACGGTATGAGGAATTGTGGCA |
| DNA template for 100-nt RNA marker. The T7 promoter is highlighted in green.<br>Transcribed region region is underlined. | TACCGAGCTCGAATTACACGAGAGCGCCATCTCCACTTTTGGAGATATCAAA<br>TTGGTAACCAGCACTGAAATTATGTGATCTGCTTTTGCGTGCAATCGGAGAA<br>TTAAAGGACCAGGTACCAGAAGCCGCGAGGACATCGAGGAGTACTGCAAA<br>CCTCTGTTAGAGTCCAATGAAGACGcgatTAATACGACTCACTATAGGATGTAT<br>GAGCATGACTACATCGGCGCCATCTTAGAGTGGGGCAAGACCAAGATCCTG<br>TCTCCGCATATCATTAAGTATCGGTATGAGGAATTGTGGCA |

| | |
|---|---|
| DNA template for 50-nt RNA marker. The T7 promoter is highlighted in green. Transcribed region region is underlined. | TACCGAGCTCGAATTACACGAGAGCGCCATCTCCACTTTTGGAGATATCAAATTGGTAACCAGCACTGAAATTATGTGATCTGCTTTTGCGTGCAATCGGAGAATTAAAGGACCAGGTACCAGAAGCCGCGAGGACATCGAGGAGTACTGCAAACCTCTGTTAGAGTCCAATGAAGACGTTAGAGTGGGGCAAGACCAAGATCCTGTCTCCGCATATCATTAAGTATCGcgatTAATACGACTCACTATAGGATGTATGAGCATGACTACATCGGCGCCATCGTATGAGGAATTGTGGCA |
| **DNA oligonucleotides** | **Sequence (5'-3')** |
| *Oligonucleotides used for structure determination* | |
| Non-target strand (60 nt) | CTAGCGTTGTGGGATACCCGTTACACTAATTTTTCGGTTCACGATATCCTTTAGTCCCAC |
| Target strand (60 nt) | GTGGGACTAAAGGATATCGTGAACCGAAAAATTAGTGTAACGGGTATCCCACAACGCTAG |
| Non-target strand (99 nt) | CTAGCGTTGTGGGATACCCGTTACACTAATTTTTCGGTTCACGATATCCTTTAGTCCCACAACTAACGACTACACTTTTTGGGTGACCGACCCAACACG |
| Target strand (99 nt) | CGTGTTGGGTCGGTCACCCAAAAAGTGTAGTCGTTAGTTGTGGGACTAAAGGATATCGTGAACCGAAAAATTAGTGTAACGGGTATCCCACAACGCTAG |
| Target strand (99 nt, mismatched at transcription bubble) | CGTGTTGGGTCGGTCACCCAAAAAGTGTAGTCGTTAGTTGTCCCTGATTTCCTATACGTGAACCGAAAAATTAGTGTAACGGGTATCCCACAACGCTAG |
| *Oligonucleotides used to introduce mutations* | |
| dCas12f S106A_F | ATCCTTgcgACCCTGAACCACGGCCTG |
| dCas12f S106A_R | CAGGGTCGCAAGGATATTCGTCGGCAT |
| dCas12f H110A_F | CTGAACgcgGGCCTGATCAGCAGCTTT |
| dCas12f H110A_R | CAGGCCCGCGTTCAGGGTGGAAAGGAT |
| dCas12f Y89A_F | ACCACCgcgCGCGTGGTGAGCGACCGT |
| dCas12f Y89A_R | CACGCGCGCGGTGGTGTTCATACGGCT |
| dCas12f_R90A_F | ACCTATgcgGTGGTGAGCGACCGTTTT |
| dCas12f_R90A_R | CACCACCGCATAGGTGGTGTTCATACG |
| dCas12f Y89A/R90A_F | ACCACCgcagcgGTGGTGAGCGACCGT |
| dCas12f Y89A/R90A_R | CACCGCTGCGGTGGTGTTCATACGGCT |
| dCas12f Q144A_F | GGTCTGgcgGGTATCTCTCGTCTGGTG |
| dCas12f Q144A_R | GATACCCGCCAGACCAAACGGGAATGC |
| dCas12f F337A _F | GAGGAGgcgGTGCTGCGTAATTGGTCC |
| dCas12f F337A _R | CAGCACCGCCTCCTCCTCTTTGGCGAT |
| dCas12f F337K _F | GAGGAGaagGTGCTGCGTAATTGGTCC |
| dCas12f F337K _R | CAGCACCTTCTCCTCCTCTTTGGCGAT |
| dCas12f WSYYE-ASAAE (342–346)_F | ggctgaaCTGATGACCAAAATCAAGTAC |

| | |
|---|---|
| dCas12f WSYYE-ASAAE (342–346)_R | gcggacgcATTACGCAGCACGAACTC |
| dCas12f lid (325–346) to (GSGSGS)x3_F | agtggtagtggtagtggtagtggtagtCTGATGACCAAAATCAAGTAC |
| dCas12f lid (325–346) to (GSGSGS)x3_R | accactaccactaccactaccactaccATTCATAAGGATCAGGGTG |
| $\sigma^E$ F225A_F | TACAGCgcgTCTGAGATTGCCAAGGAG |
| $\sigma^E$ F225A_R | CTCAGACGCGCTGTACTTCAAATCGCA |
| $\sigma^E$ H144A_F | AAGCGCgcgCTGATCGAGTTATGTCTG |
| $\sigma^E$ H144A_R | GATCAGCGCGCGCTTATCCGCATTCAG |
| $\sigma^E$ Y152A_F | CTGAAGgcgGGTTTCCAATATAAGGCG |
| $\sigma^E$ Y152A_R | GAAACCCGCCTTCAGACATAACTCGAT |
| $\sigma^E$ H168A_F | GGTATTgcgGAAACCAGCCGTGAAATC |
| $\sigma^E$ H168A_R | GGTTTCCGCAATACCTTTACCCATAAC |
| $\sigma^E$ H130A_F | TTCGACgcgATTAAATCGATTCTTCCG |
| $\sigma^E$ H130A_R | TTTAATCGCGTCGAAGGATTTCTGGGT |
| $\sigma^E$ L137A_F | CTTCCGgcgCTGAATGCGGATAAGCGC |
| $\sigma^E$ L137A_R | ATTCAGCGCCGGAAGAATCGATTTAAT |
| $\sigma^E$ I184A_F | AAGACCgcgGTCCACAAGGGCAACGAG |
| $\sigma^E$ I184A_R | GTGGACCGCGGTCTTGATGTCCTCGAT |
| $\sigma^E$ Y105A_F | CAGGACgcgATGGTGGGTTATGATCCG |
| $\sigma^E$ Y105A_R | CACCATCGCGTCCTGATAGTTGTCGTA |
| $\sigma^E$ F72A_F | TTGCGTgcgGTGATGAAGCGCGAGTGC |
| $\sigma^E$ F72A_R | CATCACCGCACGCAAGAAGTAAAAAAT |
| $\sigma^E$ R76A_F | ATGAAGgcgGAGTGCATTAGCTATTAC |
| $\sigma^E$ R76A_R | GCACTCCGCCTTCATCACGAAACGCAA |
| $\sigma^E$ F72A/R76A_F | gaaggccGAGTGCATTAGCTATTAC |
| $\sigma^E$ F72A/R76A_R | atcaccgcACGCAAGAAGTAAAAAATG |
| $\sigma^E$ H240A/F243A_F | gaggccATGAAGGCATATAAACTGATG |
| $\sigma^E$ H240A/F243A_R | ttgcgcTACCTCTTTCTCGGACAG |
| $\sigma^E$ H65A_F | CCGGAAgcgATTTTTTACTTCTTGCGT |
| $\sigma^E$ H65A_R | AAAAATCGCTTCCGGGCTTTCCAGCTT |
| $\sigma^E$ W35A_F | CGTCAGgcgCTGAACGATGATTTTTTG |
| $\sigma^E$ W35A_R | GTTCAGCGCCTGACGGCCGATCCACAG |
| $\sigma^E$ Y82A_F | AGCTATgcgCGTAAACCAAAATACAAA |
| $\sigma^E$ Y82A_R | TTTACGCGCATAGCTAATGCACTCGCG |
| $\sigma^E$ R172A_F | ACCAGCgcgGAAATCAAAGAAGCCATC |

| | |
|---|---|
| $\sigma^E$ R172A_R | GGTATTCACGAAACCAGCGCGGAAATC |
| $\sigma^E$ R83A_F | TATTACgcgAAACCAAAATACAAATTT |
| $\sigma^E$ R83A_R | GGTTTCGCGTAATAGCTAATGCACTCG |
| $\sigma^E$ R218A_F | AAGATGgcgTGCGATTTGAAGTACAGC |
| $\sigma^E$ R218A_R | ATCGCACGCCATCTTCAGCACCTTAGC |
| $\sigma^E$ L137K_F | CTTCCGaagCTGAATGCGGATAAGCGC |
| $\sigma^E$ L137K_R | ATTCAGCTTCGGAAGAATCGATTTAAT |
| $\sigma^E$ Del 201–265_F | CCAATGAATAATAAGAATTCCTGCTGCCACCG |
| $\sigma^E$ Del 201–265_R | CTTATTATTCATTGGTCATGGTGTCATTGC |
| *Oligonucleotides used for cloning* | |
| dCas12f_F | TTGTATTTCCAGGGCATGGGAAAATCAACACTAAAGCAC |
| dCas12f_R | CAAGCTTCGTCATCATTAGCCGATAATCAATTCGATGCC |
| $\sigma^E$_F | TTGTATTTCCAGGGCATGTTATATTCAGATTTTG |
| $\sigma^E$_R | CAAGCTTCGTCATCATTACGCGCTTTGCAGTTGC |
| dCas12f/$\sigma^E$_F | TTGTATTTCCAGGGCATGGGAAAATCAACACTAAAGCAC |
| dCas12f/$\sigma^E$_R | CAAGCTTCGTCATCATTACGCGCTTTGCAGTTGC |
| $\sigma^E$ Vector Del His_F | ATGTTATATTCAGATTTTGAATTGC |
| $\sigma^E$ Vector Del His_R | TTTTTTCATGGTATATCTCCTTC |
| *rpoA_rpoB*_F | CACAGCCAGGATCCGATGGCTCTATTAAATTTTCAAAAAC |
| *rpoA_rpoB*_R | CGCCGAGCTCGAATTTTACTCTTCCAGGCGGATG |
| *rpoC_rpoZ* | Whole plasmid synthesized by Genscript |
